# Hunger modulates behavioral responses to olfactory and chemotactile cues in the specialist predator of dangerous prey, *Berghia stephanieae*

**DOI:** 10.64898/2026.05.19.726230

**Authors:** Kate Otter, Kassidy Ye, Rylie Costello, Jacqueline Forbes, A. Cairo Laurenzcia, Paul S. Katz

## Abstract

Animals continuously evaluate environmental cues to guide approach-avoidance decisions, with internal states like hunger dynamically shaping how stimuli are acted upon. While most studies examine the valence-switching of stimuli from appetitive to aversive using simplified or ambiguous stimuli, we leveraged a system in which a single prey contains both appetitive and aversive features. The nudibranch *Berghia stephanieae*, is a specialist predator of the sea anemone, *Exaiptasia diaphana*. These nudibranchs must resolve conflicting signals where chemical cues signal food, while contact can result in injury or death. The danger posed by *Exaiptasia* was described and quantified through direct counts of nematocysts fired into *Berghia* and multiple instances where the *Berghia* was captured and consumed by its prey. To test how internal state influenced the perception of stimuli from prey we recorded predatory behavior of *Berghia* after different periods of food deprivation. We found that the olfactory cues from prey were attractive to *Berghia*, even when animals were sated, and usually led to a contact-mediated investigation of prey. Hunger independently modulated olfactory and contact cue valence at different internal states and time scales of food deprivation. Hunger specifically altered the threshold for avoidance following contact with prey, indicating that somatosensory and chemotactile cues are modulated by hunger unlike olfactory cues. Our results highlight how internal state and sensory modality interact to shape decision making in a biologically relevant, high-risk predation context.

## Introduction

Animals evaluate sensory information in the context of internal states to generate appropriate behavioral responses. Many stimuli can be either attractive or aversive depending on an animal’s internal state, with valence-switching allowing for adaptive decision making (Hirayama and Gillette, 2012; Tye, 2018; Rengarajan *et al*., 2019; Kanwal and Parker, 2022). Hunger is one of the most powerful states driving animal behavior (Smith and Grueter, 2022); it can promote risk-taking (Moran *et al*., 2021) and reduce sensitivity to aversive stimuli (Gaudry and Kristan, 2009; Inagaki, Panse and Anderson, 2014). In diverse species, food deprivation can reverse natural aversions, as seen in *Drosophila melanogaster* larvae shifting responses to odorants, adult flies modulating CO_2_ avoidance based on foraging state, and *Caenorhabditis elegans* becoming attracted to CO_2_ after starvation (van Breugel, Huda and Dickinson, 2018; Rengarajan *et al*., 2019; Vogt *et al*., 2021). In predatory animals, prey stimuli may be perceived as appetitive when hungry but aversive or neutral when sated. For example, the predatory gastropod *Pleurobranchaea californica* exhibits bidirectional valence switching, responding appetitively to noxious stimuli when hungry and aversively to appetitive stimuli when sated (Gillette *et al*., 2000). This dynamic is particularly relevant when prey are dangerous, such as toxic or defensive species, where appetitive and aversive cues may overlap, providing a natural context to study how satiety influences cue perception.

Here, we are investigating the feeding ethology of the nudibranch mollusc, *Berghia stephanieae*. Like many nudibranchs, *Berghia is* a monophagous predator specializing on the sea anemone *Exaiptasia diaphana* (Goodheart *et al*., 2017; Monteiro *et al*., 2020). *Exaiptasia* defends itself with nematocysts and acontia—thread-like structures densely packed with nematocysts and specialized anti-predator toxins (Rodríguez *et al*., 2012; Lam *et al*., 2017; Ashwood *et al*., 2022). Given their specialization, it was long thought that nudibranchs experience minimal risk of death due to defenses such as nematocyst sequestration (Goodheart and Bely, 2017), mucosal defenses (Greenwood *et al*., 2004), and specialized behavior (Edmunds *et al*., 1976; Conklin and Mariscal, 1977; Seavy and Muller-Parker, 2002; Garese *et al*., 2012). However, recent evidence has shown that cnidarians harm and even feed upon their sea slug predators in the field (Mehrotra *et al*., 2019; Hayes and Schultz, 2022).

We hypothesized that hunger increases *Berghia’s* attraction to and interaction with prey by shifting the perceived valence of stimuli from prey from aversive to attractive. The impact of hunger on predatory behavior was assessed by recording *Berghia* in an arena with *Exaiptasia* and using a combination of manual event logging software and automated tracking to quantify behavioral interactions. We developed a detail ethogram of predatory behavior in *Berghia* and in Experiment 1, we validated that *Exaiptasia* are dangerous prey that injure and can consume *Berghia*. We predicted that due to the danger of their prey, sated *Berghia* would respond aversively to prey cues and that hunger would cause animals to be attracted to their prey and have appetitive responses to their prey. Contrary to our hypothesis, *Berghia* are always attracted to their prey, while hunger shifts the threshold for avoidance in response to contact-mediated cues.

## Results

### *Berghia* sustained injury from their prey during hunting, handling and feeding

Individual *Berghia* were observed being captured by their prey several times in the lab (Figure 1 A). This typically happened when the anemone had adhered itself to the substrate, which allowed the anemone to bend its body column after wrapping tentacles around the *Berghia*. However, when detached from the substrate, the anemones usually were unable to lift the *Berghia* onto their oral disc for ingestion (see Figure 1 B for counter example). Therefore, to minimize the risk of slugs being captured during the experiments that involved repeated measures within animals, anemones were not allowed to attach to the substrate of the testing arenas.

**Figure 1.**
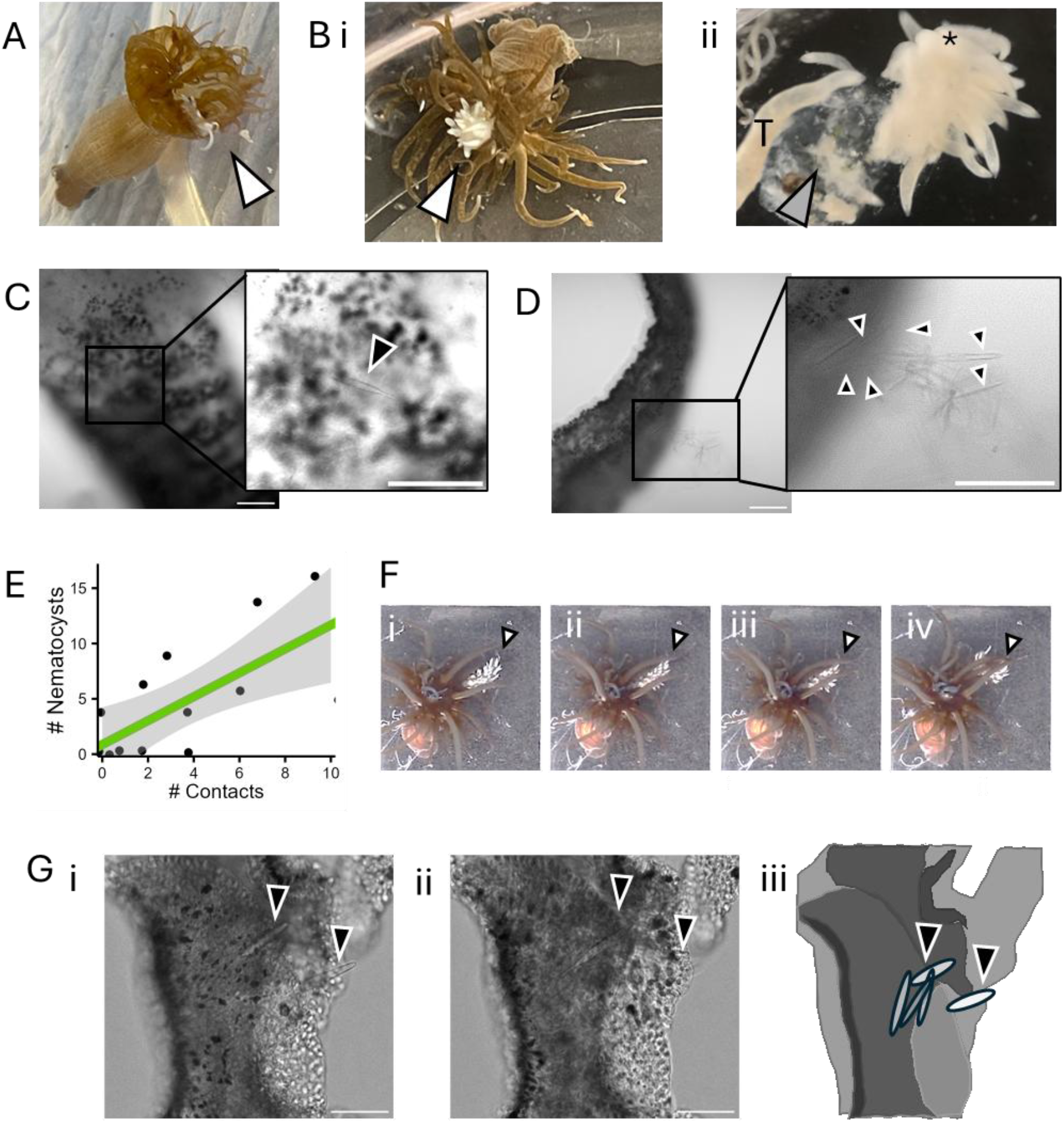
Berghia is stung by nematocysts during contact with its anemone prey. A) Berghia was captured by anemones most often when the anemone is securely attached to the substrate. This image was captured after a spontaneous occurrence of a Berghia capture. The white arrow indicates the Berghia’s head. B) It is rarer that a Berghia is caught by an anemone that is not securely attached to the substrate. i) In this example, the slug attempted to struggle free and was able to free its head (indicated by the black arrow) in about two minutes after being captured. ii) When the same slug was removed from the anemone with forceps, the posterior half of the body (indicated by the gray arrow), was digested by the anemone (tentacle indicated by the “T”, head of the slug indicated by *). C) An oral tentacle that was removed from a slug within five minutes of interacting with an anemone. It was fixed and mounted on a slide in 100% glycerol. DIC confocal imaging revealed nematocysts embedded in the tissue, as indicated by the black arrow in the inset (black box). D) In some instances, there were many fired nematocysts (black arrows) in one area of an oral tentacle (visible in the black box outlined inset). E) The number of nematocysts counted in the face of each Berghia was positively correlated with the number of contacts between the slug and the anemone. The linear regression line (green) with a slope of 1.08 was statistically significant (F(1, 12) = 11, p = 0.006), with an adjusted R² of 0.435.F) Consecutive images of an interaction between Berghia and E. diaphana with a black arrow indicating the tail of the Berghia. From left to right: i. the slug is feeding on an anemone, ii. the anemone moves its tentacles towards the tail of the slug, iii. the anemone makes contact with the tail, and iv. the slug stops feeding and turns away from the anemone. G) Confocal images of the tail of the same slug depicted in F. From left to right: i. and ii. two slices from the confocal stack of the tail from F and G, that show the nematocysts embedded in the tail (indicated by the black arrows), iii. a schematic of the tail tissue with nematocysts indicated by the black arrows. All scale bars are 100 μm in length.

There were numerous nematocysts embedded in the skin of the slug following interactions (Figure 1 C, D). The number of embedded nematocysts was positively correlated with the number of times the anemone made contact with the *Berghia* (Figure 2E; F(1, 12) = 11, p = 0.006, R² = 0.435). The regression coefficient for the number of contacts was 1.08, with a standard error of 0.033, suggesting that, on average, each contact resulted in an increase of approximately one nematocyst on the oral tentacles or lips of the *Berghia*. It is likely that the number of fired nematocysts represents an underestimate due to potential loss during dissection, fixation, and mounting of the tissue.

**Figure 2.**
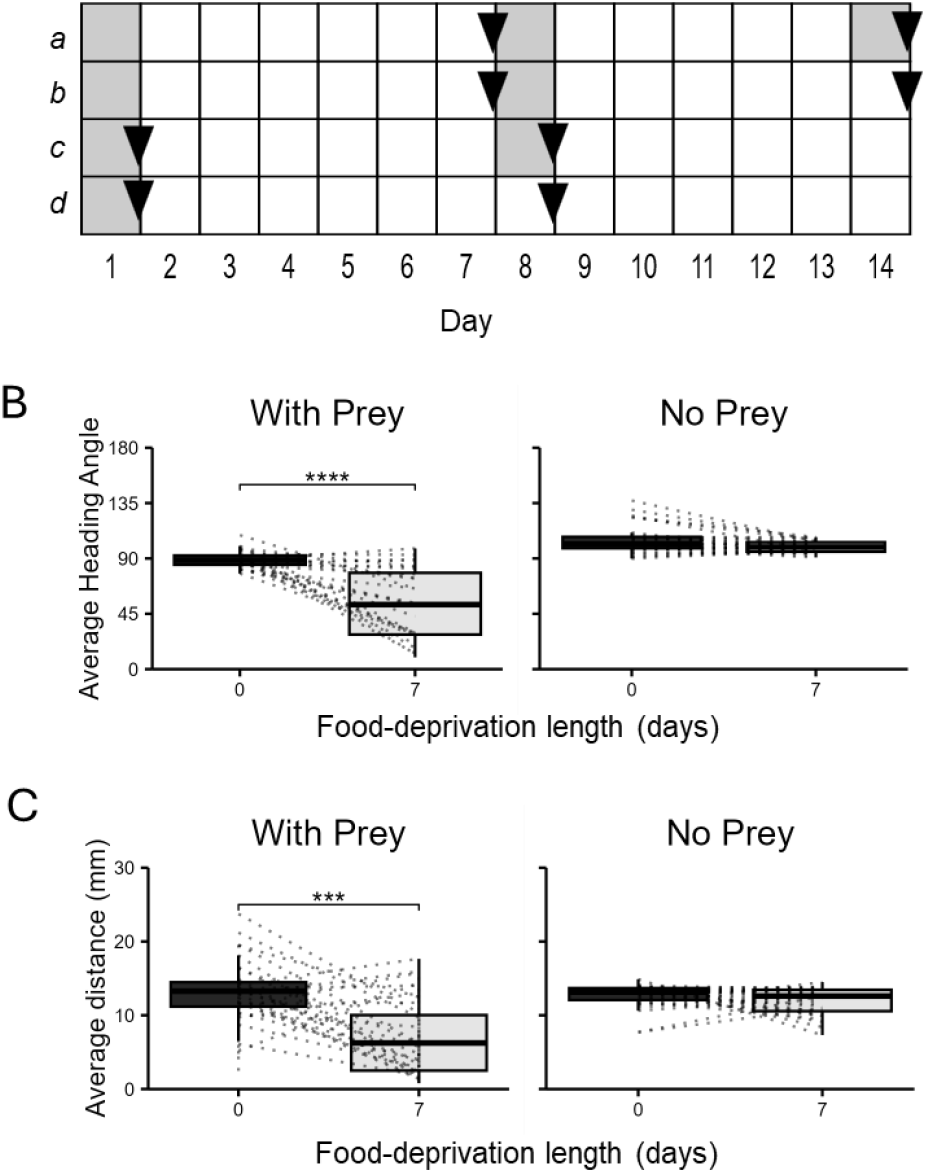
Food deprivation and presence of prey drive changes in behavior. Food deprivation and presence of prey drive changes in behavior. A) Schematic representing the food deprivation regimes used in this study. Each box represents one 24-hour period. The grey boxes represent ad libitum access to anemones. The black triangles represent when the experiments were conducted. The groups included slugs that were: (a) first tested after 7-days of food deprivation and then tested sated, (b) 7-days food-deprived both times they were tested, (c) sated both times they were tested, and (d) slugs that were tested sated and after 7-days of food deprivation. B) The average heading angle from prey in degrees for 0- and 7-day food-deprived animals. The average heading angle is calculated from the center of the oral disc of Exaiptasia, the middle of the slug’s body, and their face. Dotted lines connect trials from an individual animal. Left is slugs in an arena with their prey and right is slugs in an empty arena. Asterisks show significant relationships as tested by a paired t-test (t(25) = 4.561, p = 1.16e-4, Cohen’s d = 0.894). C) Plots arranged as in B representing the average distance from prey in mm. Asterisks show significant relationships as tested by a paired t-test (t(25) = 6.112, p = 2.18e-6, Cohen’s d = 1.199).

In some instances, behavioral interactions could be directly related to the nematocysts found embedded in the *Berghia*. For example, after a touch to the tail that caused the *Berghia* to discontinue feeding and turn away from its prey (Figure 1 F), imaging of the tail revealed five nematocysts embedded in it (Figure 1 G, H).

### *Berghia* exhibited diverse, stereotyped predatory behaviors

To quantify predatory behavior in *Berghia*, we developed an ethogram based on detailed observations of individuals in uniformly lit arenas (Table 1, Video 1). Behaviors were described from approximately 50 individuals. Over continued laboratory observations and data analysis, the descriptions of each behavior were clearly defined, and additional behaviors were added as needed. The described ethogram based on these observations was used for this study. Many of these behaviors consisted of discrete, stereotyped movements. The ethogram, which ultimately included about 20 behaviors (Table 1), was developed collaboratively by five lab members, with each behavior corroborated by at least two observers before inclusion. After finalizing the ethogram, we compared our results to previously published ethograms of other nudipleuran species, identifying potential synonymous behaviors (Supplemental Table 1).

**Table 1.**
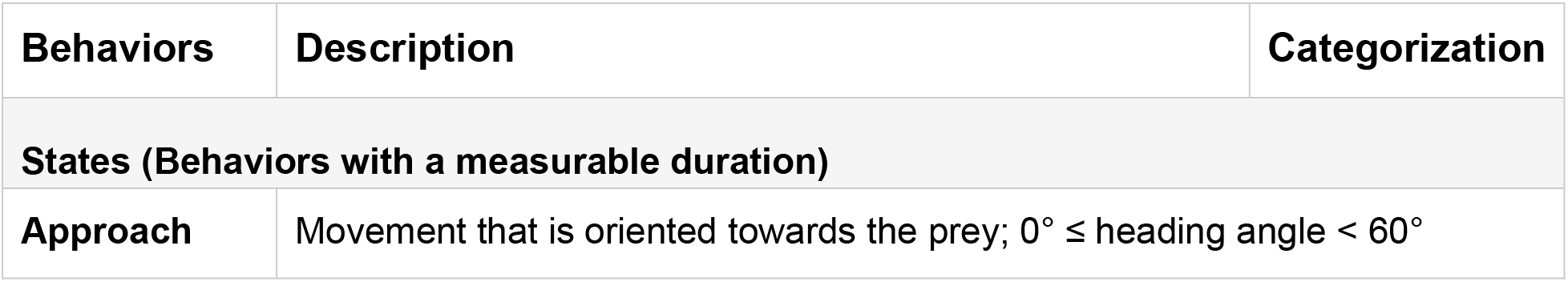
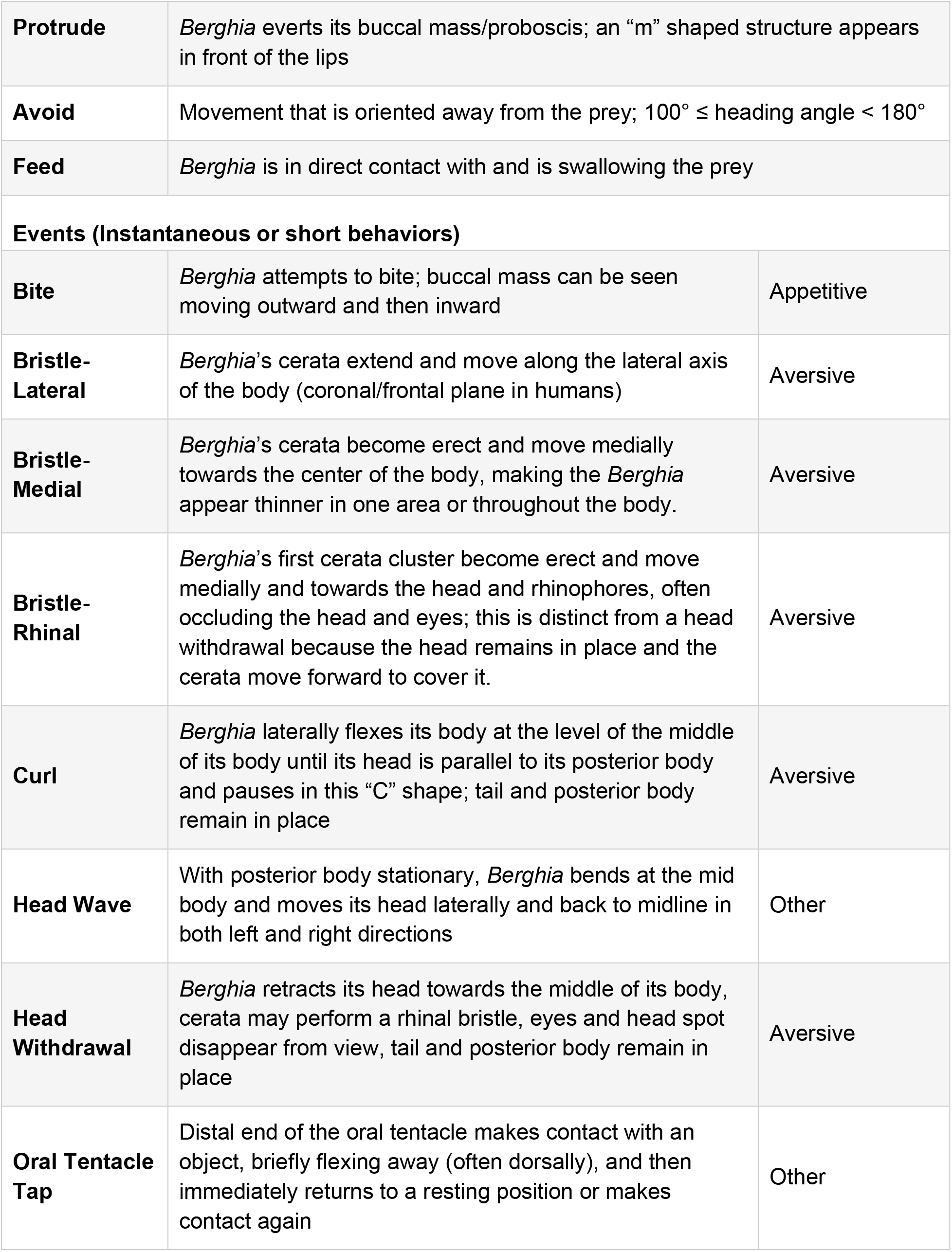
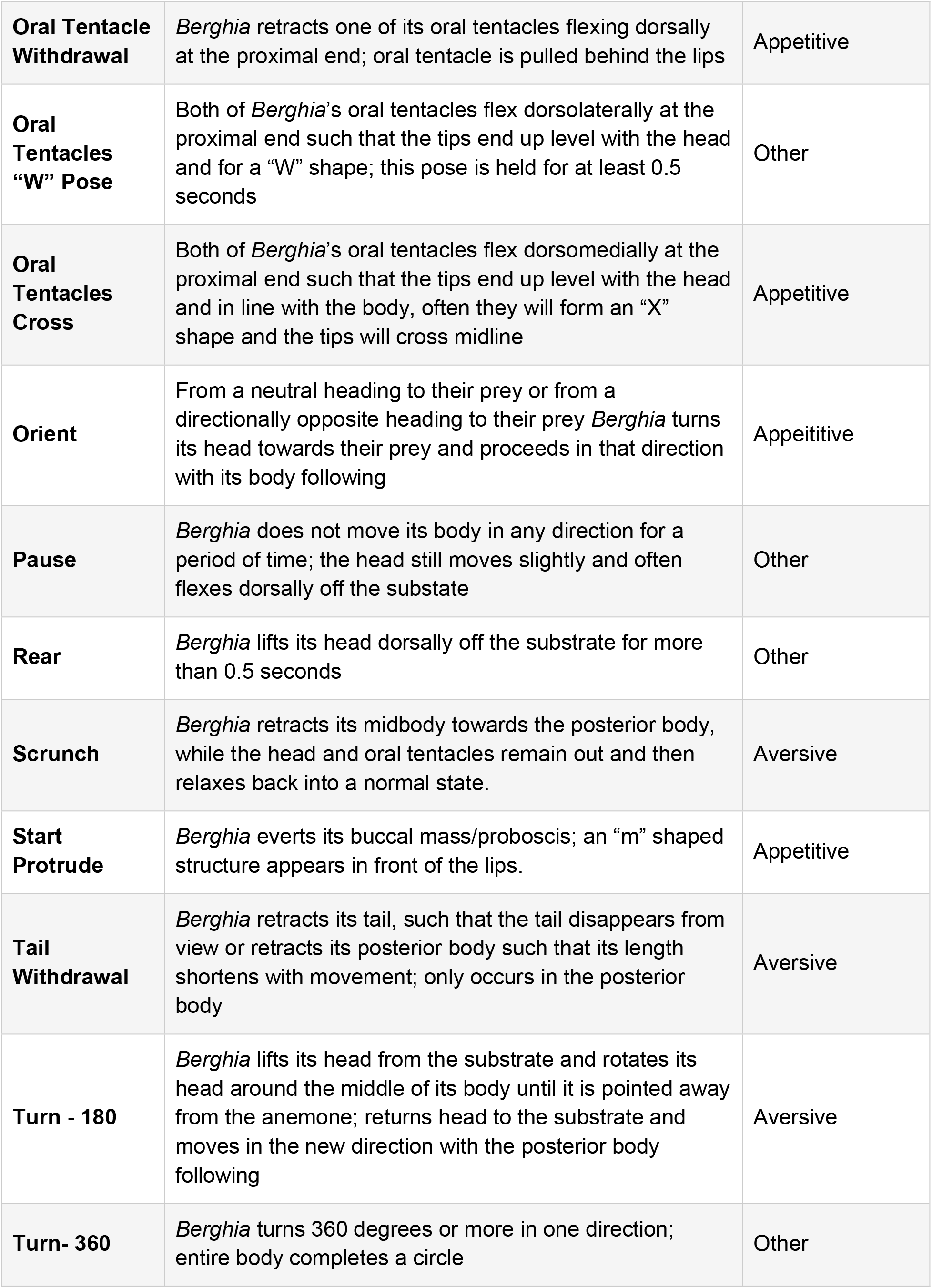
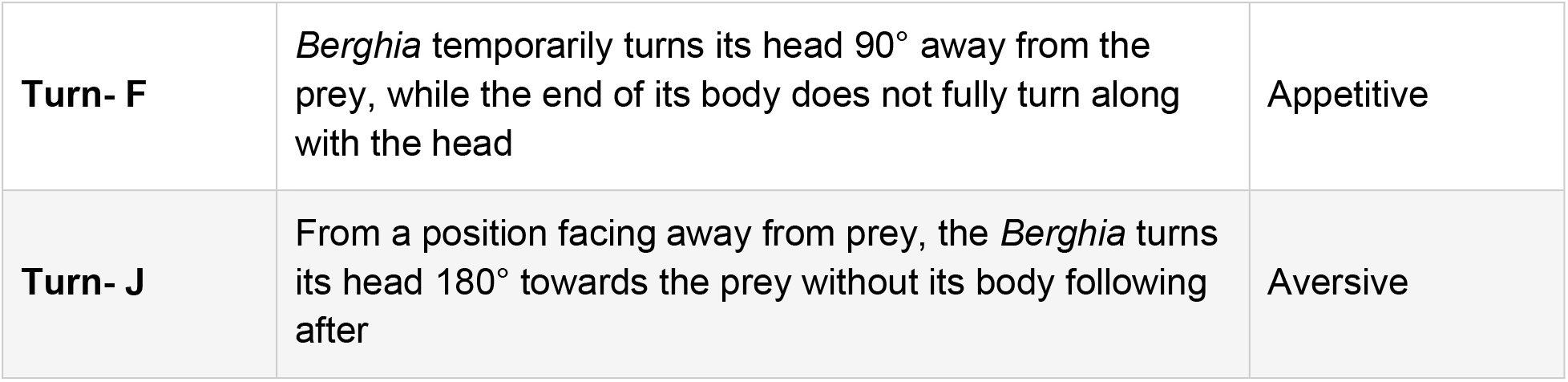
Ethogram of foraging and predatory behaviors characterized in Berghia.

For the purposes of this study, we categorized behaviors into three categories: appetitive, aversive and other. Appetitive behaviors were those that were clearly associated with feeding (e.g., Cross Oral Tentacles or Protrude) or that decreased the distance between the slug and their prey (e.g., orient, J-turn). Aversive behaviors were those that increased distance between the slug and their prey (e.g., tail withdrawal, 180-Turn). Bristling of the cerata was also considered a behavior that indicated that the animal perceived a stimulus as aversive, in accordance with other studies (Brown *et al*., 2024). Other behaviors were those that did not clearly fall into either category.

To validate the intuitive categorization of behaviors, we identified the probability that each behavior occurred within 15 seconds of feeding (appetitive behaviors) or within 15 seconds of turning away from their prey (aversive behaviors). This validated the categorization of most behaviors that were not already known to be appetitive or aversive (e.g., bristling, tail withdrawal). One notable exception was the oral tentacle withdrawal, which does increase distance between *Berghia* and their prey. However, it moves a single oral tentacle out of the way of the mouth, much like the cross oral tentacles behavior, which moves both oral tentacles.

The most common behavior when a *Berghia* interacted with any object, including prey, was an oral tentacle tap. This behavior was characterized by a brief touch with the tip of the oral tentacle then a bend near the tip away from the object touched with slight retraction of the oral tentacle, then immediately returning to the original position and often contact with the object or surface (Table 1). This behavior was not clearly appetitive or aversive, because it occurred during contact with most objects and only briefly increased the distance between the oral tentacle tip and object before returning to contact. When touching anemones, several taps often led to a withdrawal (aversive) or cross oral tentacles posture (appetitive). Thus, this behavior is likely exploratory in nature and therefore was not classified as an appetitive or aversive reaction in this study.

### Hunger drove changes in behavior in the presence of prey

We first quantified the behavior of *Berghia* when they were 0- and 7-day food-deprived and placed in an arena with their prey (Figure 2 A). Unsurprisingly, animals spend more time feeding when they are 7-day food-deprived (M = 228.471 sec, SD = 241.243), than when they are sated (M = 0, SD = 0), (t(25) = 6.112, p = 2.18e-6, Cohen’s d = −0.947). The average heading angle, which indicates the degree to which they were oriented toward their prey, was smaller when the slugs were 7-days food-deprived (M = 54.261 degrees, SD = 28.158) than when they were sated (Figure 2 B; M = 89.589, SD = 7.079), (t(25) = 6.112, p = 2.18e-6, Cohen’s d = 1.199). The presence of prey reduced the heading angle of both sated and food-deprived animals. The average heading angle to the center of the arena when sated in an empty arena (M = 105.128, SD = 11.52) was significantly larger than sated animals in the presence of prey (Figure 2E; t(51)= 6.388, p = 4.98e-8, Cohen’s d = 1.656). Food-deprived animals in an empty arena (M = 99.17, SD = 4.674) also had significantly larger heading angles than 7-day food-deprived animals in the presence of prey (Figure 2 B; t(28) = 8.575, p = 2.37e-9, Cohen’s d = 2.289). There was no effect of repeated testing (Supplementary Figure 3) on any of the metrics measured.

We found that *Berghia* spent more time close to their prey when they were 7-day food-deprived (M = 7.449mm, SD = 5.361) than when they were sated (M = 23.589 mm, SD = 4.374), (Figure 2 C; t(25) = 4.561, p = 1.16e-4, Cohen’s d = 0.894). The prey was generally in the center of the arena and slugs often spent their time circling the edge of an empty arena. To test whether the slugs exhibited hunger induced hyperactivity that lead to spending more time in the center of the arena, we also recorded the distance and heading angle from the center of an empty arena for 0- and 7-days food deprived slugs. The changes in distance were specific to arenas containing prey for 7-day food deprived animals. The distance to a fixed point in 7-day food-deprived animals in an empty arena (M = 11.895, SD = 1.837) was significantly greater than the mean distance to the prey (t(33) = 4.546, p = 7.11e-5, Cohen’s d = 1.209).

To test whether hunger induces exploratory behavior for any object in an arena, slugs were tested in an arena with an opaque white marble in the center. These behavioral changes were specific to an arena containing prey; in an arena with a white marble placed in the center, they behaved similarly to an empty arena when sated and food-deprived (Supplemental Figure 4). Other features of their trajectories within the arenas were analyzed including straightness, distance traveled, directional change, sinuosity, mean speed and standard deviation of speed and directional change (Supplemental Table 3). These trajectory features were compared using two-way ANOVAs to investigate the effect of experiment type, food deprivation and the interaction between food deprivation and experiment type. There was no interaction between food deprivation and the experiment type (empty arena, arena with prey, arena with marble) for any of the metrics measured (Supplemental Table 3). *Berghia* in prey-containing arenas traveled less distance (F(2,185) = 9.017, p = 0.0038; Supplemental Figure 5) and showed more directional change (F(2,185) = 7.839, p = 0.011; Supplemental Figure 5).

### Food deprivation gradually changed proximity and orientation to prey, resulting in increased feeding

For this experiment, rather than using a hungry/sated binary, we used progressive lengths of food-deprivation to understand how hunger is modulating decision making. We tested 60 individual animals across 0 to 5 days of food deprivation, with animals included in the analysis only if they completed at least 4 trials. To control for order effects, animals were divided into 6 groups that experienced the food deprivation lengths (FDLs) in different orders, resulting in a total of 343 trials analyzed (Figure 3). Food deprivation decreased the average heading angle gradually with a plateau around 3 days of food deprivation (Figure 3 B; GLMM; B = −0.11, SE = 0.015, t = −7.41, p =1.28e-13). Food deprivation also gradually decreased the mean distance from prey (Figure 3 C; GLMM; B = −0.12, SE = 0.016, t = −7.39, p = 1.5e-13) and duration of time avoiding (Figure 3 D; GLMM; B = −0.13, SE = 0.035, z = −3.73, p = 0.00019). That said, sated animals spent only about an average of 73 seconds avoiding which was less than 10% of the trial duration. Unlike the other measures of predatory behavior, the duration of time spent feeding increased gradually and did not level off after 3-days (Figure 3 E; GLMM; B = 0.23, SE = 0.036, z = 6.33, p = 2.47e-10).

**Figure 3.**
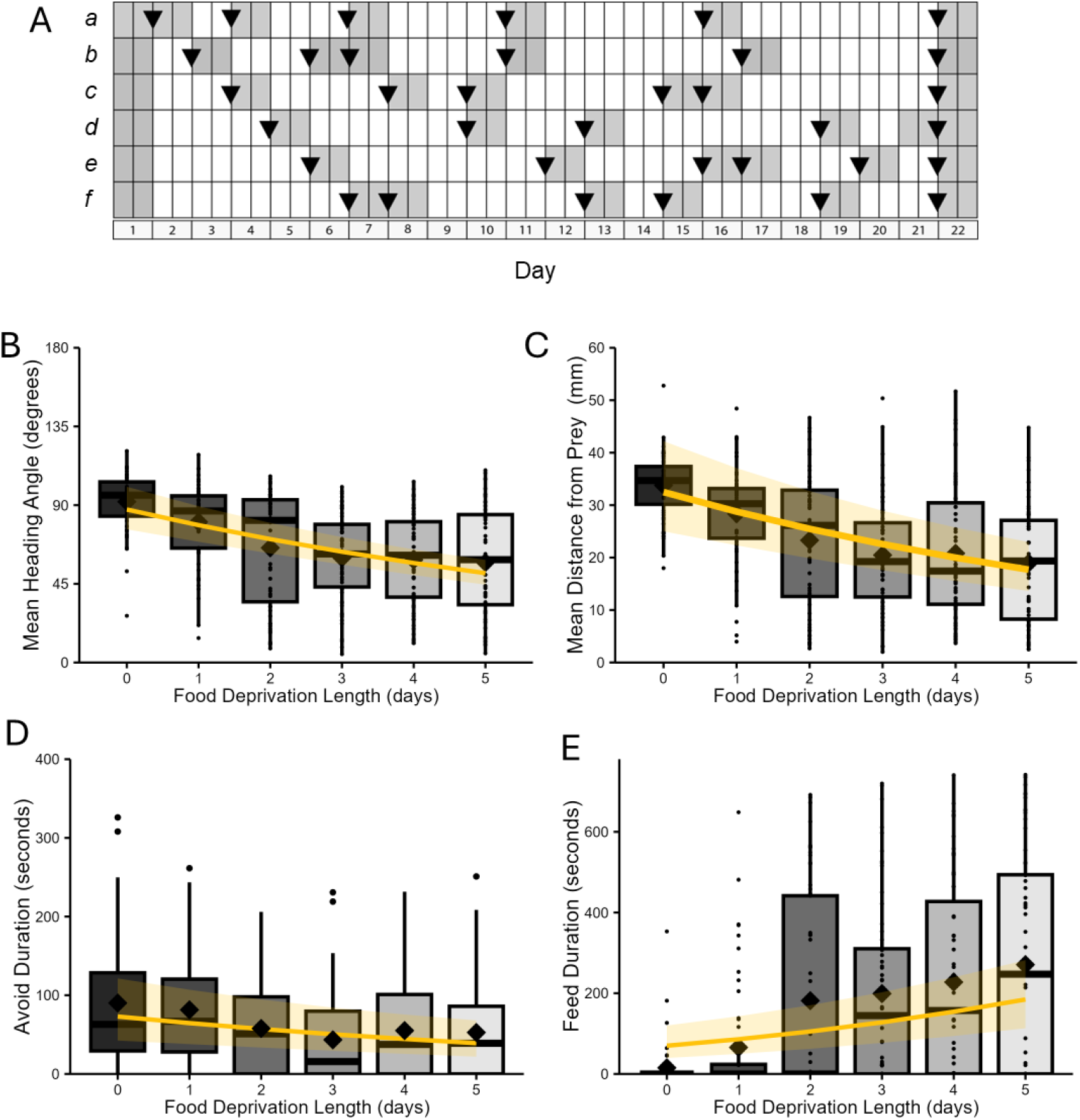
Food deprivation gradually changes behavior in the presence of prey. A) The food deprivation regimes used in this study. Each box is a 24-hour period, grey boxes represent 24-hour access to prey, black triangles represent trials. Each row (a-f) represents a group of about 10 animals that underwent that feeding and trial regime. B-E) The yellow line represents the fixed effects from a fitted mixed effects model with the standard error represented by the lighter yellow shading around the line. Diamonds represent the mean. For all mixed effects models animal identity nested within the experiment year were included as random intercepts. B) Average heading angle in degrees for each FDL. The fitted model used a gamma family with a log link function. C) Average distance from prey in mm throughout the trial. The fitted model gamma family with a log link function. D) The duration of time Berghia spent avoiding its prey. The fitted model used a beta family, where each duration of time was divided by the total time (780 seconds) and then we added a small constant (0.001) to allow the values to be within 0 and 1. Values were back-transformed for the plot. E) Duration of time feeding (seconds) for each FDL. The fitted model used a beta family, where each duration of time was divided by the total time (780 seconds) and then we added a small constant (0.001) to allow the values to be within 0 and 1. Values were back-transformed for the plot.

### Olfactory cues from prey were attractants regardless of food deprivation and led to contact with prey

While food deprivation changed space utilization and increased feeding, it did not impact the number of approach bouts an animal exhibited (Figure 4 A; GLMM; B = −0.02, SE = 0.019, z = −1.13, p = 0.258). Food deprivation weakly increased the average duration of each approach bout (Figure 4 B; GLMM; B = 0.055, SE = 0.026, z = 2.16, p = 0.031) and decreased the time between approach bouts (Figure 4 C; GLMM; B = −0.056, SE =0.024, z = −2.37, p = 0.018). This indicates that although the number of approach bouts did not change with food deprivation, food-deprived animals spend slightly longer in each approach bout and their approach bouts were closer together in time.

**Figure 4.**
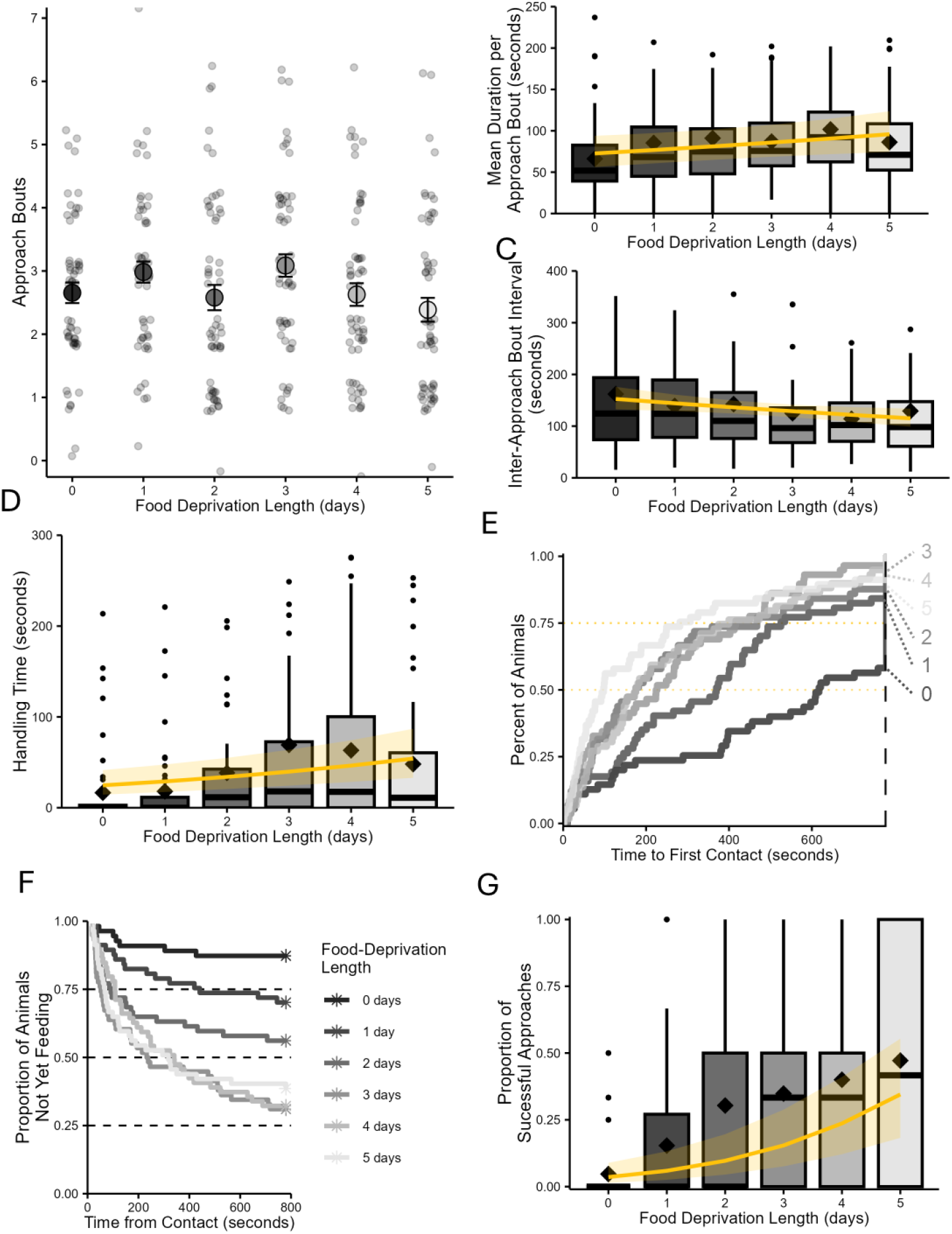
Prey are always initially attractive to Berghia, regardless of food deprivation. A) Plot showing the number of approach bouts per trial across FDL. Points are jittered to allow easier visualization. Open shaded circles represent the mean and the error bars represent the standard error of the mean. B) Boxplots showing the mean duration of each approach bout in seconds. The diamond represents the mean for each FDL. The yellow line represents the fixed effect from the fitted mixed effects model. The lighter yellow shading around the line represents the standard error. The fitted model used a gamma family with a log link function. Animal identity nested by the experiment year are included in the model as random intercepts. C) Boxplots showing the average time between approach bouts across FDL. The yellow line represents the fixed effect from the fitted mixed effects model. The lighter yellow shading around the line represents the standard error. The fitted model used a gamma family with a log link function. Animal identity nested by the experiment year are included in the model as random intercepts. D) Amount of time the Berghia spent handling their prey across FDL. The yellow line represents the fixed effect from the fitted mixed effects model. The lighter yellow shading around the line represents the standard error. The fitted model used a beta family with a logit link function, where each duration of time was divided by the total time (780 seconds) and then we added a small constant (0.001) to allow the values to be within 0 and 1. Animal identity nested by the experiment year are included in the model as random intercepts. E) Cumulative empirical density plot showing percent of animals that make contact with the anemone over time. Dotted yellow lines show 50% and 75%. The FDL is indicated by the color of the line and the number in days located on the right of the plot. Each number is connected to the end of the corresponding line by a dotted line. F) A survival plot that shows the time to feed following contact. A mixed effects Cox survival analysis was fit to the data. G) Plot showing the proportion of approach bouts that resulted in feeding across FDL with a yellow line representing the fixed effect from the fitted mixed effects model. The lighter yellow shading around the line represents the standard error. The fitted model used a binomial family. Animal identity nested by the experiment year are included in the model as random intercepts.

The time spent handling their prey also increased with food deprivation (Figure 4 D; GLMM; B = 0.16, SE = 0.31, z = 5.15, p = 2.56e-07), indicating that during these longer approach bouts the animals were within the distance of contact from the anemone. About 58% (32/55) of sated animals made contact with their prey. Over 80% of 1- and 2-day food-deprived animals made contact with their prey (48/57 and 50/57 respectively) and over 90% of 3-, 4- and 5-day food-deprived animals made contact with their prey (Figure 4 E; 56/58, 56/59 and 52/57 respectively).

Increasing food deprivation significantly accelerated the likelihood of feeding following contact with prey (Figure 4 F; mixed-effects Cox model; B = 0.35, SE = 0.048, z = 7.24, p < 0.0001, HR = 1.41). This reflects a 41% increase in the instantaneous probability of feeding at any given moment following contact, indicating that more food-deprived animals were quicker to feed. The model included random intercepts for Animal ID nested within Experiment Year to account for repeated measures. The difference in behavior was not how much they approach, but rather the outcome of these approach bouts. Ultimately, food deprivation increased the proportion of approach bouts that resulted in feeding (Figure 4 G; GLMM; B=0.53, SE = 0.10, z = 5.30, p = 1.18e-07). This indicates that contact cues are mediating decisions to approach or avoid after initial attraction to the anemone.

### Hunger shifted the avoidance threshold for contact with prey

To determine how food deprivation influences *Berghia*’s behavioral responses to contact with the anemone, we identified the behavior(s) following each type of contact and calculated how many responses to each type of contact were appetitive or aversive (see Table 1). Then we took the average response and rounded it to 0 or 1 to represent whether the net response to that contact was appetitive or aversive. We divided the slug’s body into six regions: body, head, lips, proboscis, proximal oral tentacle and distal oral tentacle. Using the total number of each type of contact per trial, the weighted average proportion of appetitive responses was calculated.

The only significant predictor of the type of response to contact with the anemone was the part of the slug’s body of the contact. Aversive responses were more likely than appetitive responses after the anemone made contact with the body (GLMM; B=-3.44, SE=0.65, z=-5.27, p= 1.35e-07) and lips (Figure 6 A; GLMM; B=1.87, SE=0.57, z=3.25, p= 0.00116). Appetitive responses were more likely in response to contact with the proboscis (GLMM; B=5.27, SE=0.50, z=10.65, p<2e-16), proximal oral tentacle (GLMM; B=3.96, SE=0.49, z=8.12 p=4.87e-16) and distal oral tentacle (Figure 6 A, B; GLMM; B=3.47, SE=0.78, z=7.25, p=4.32e-13). The parts of the body that often resulted in aversive responses were generally parts of the body that were touched by the anemone, rather than the proboscis and oral tentacles which often initiated contact.

FDL did not predict the type of response to contact neither aggregated across body-parts nor as an interaction term with contact location (Figure 6 C) and was therefore omitted from the final model. However, FDL did affect the actions that *Berghia* took. When *Berghia* was in close range of prey, it used the tips of its oral tentacles to briefly tap the space in front of it (“oral tentacle taps”, OT-taps, Table 0). These behaviors appeared to be exploratory in nature, occurring when the animal was faced with an object (e.g. a marble), conspecifics, or the edges of an arena. Food deprivation increased the total number of OT-taps (Figure 5 D; GLMM; B = 0.10, SE = 0.015, z = 6.75, p = 1.51e-11).

**Figure 5.**
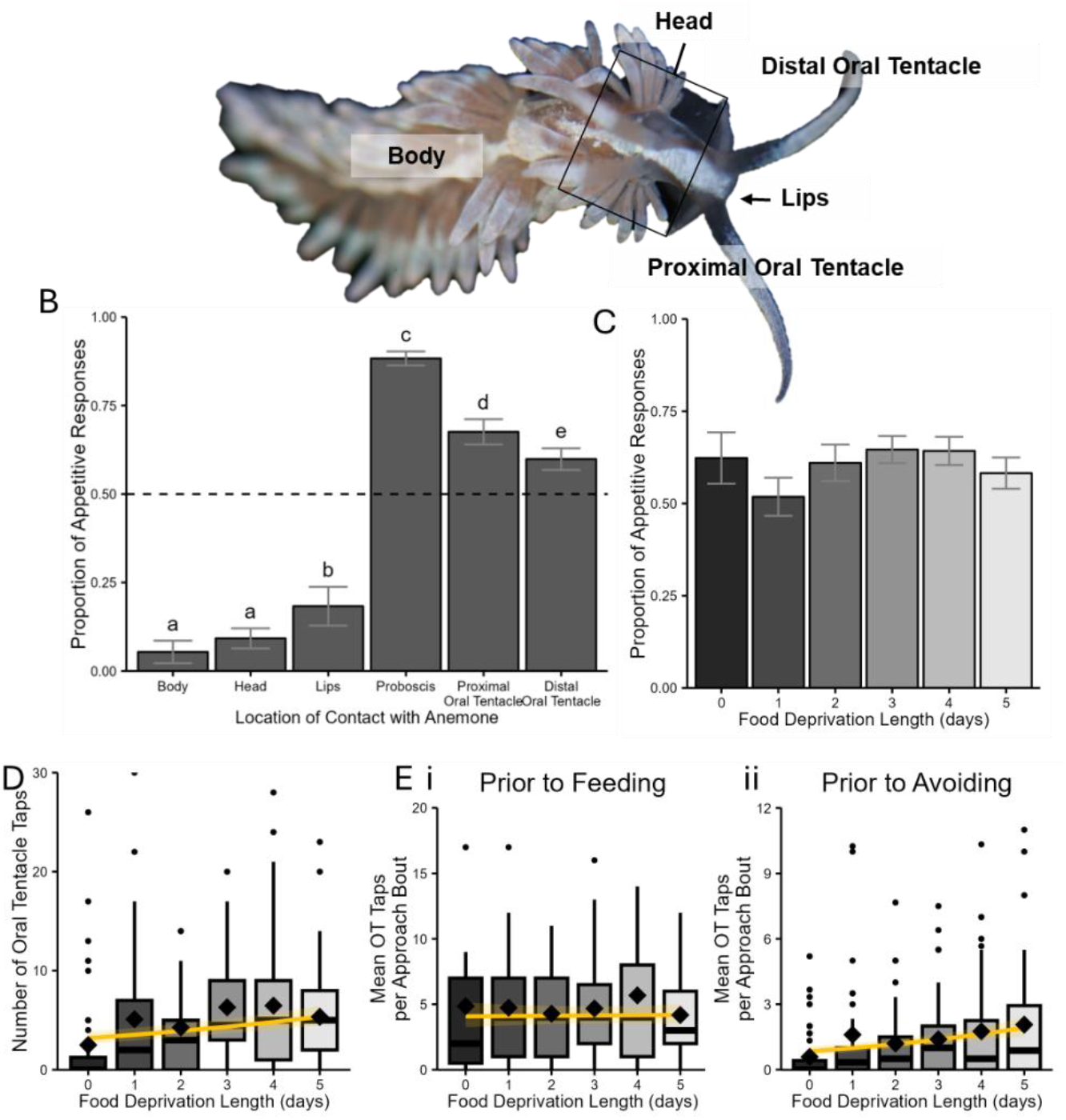
Location of contact, not FDL drives differences in response to contact while food deprivation increases sampling events. A) Image of a Berghia showing the body parts/body regions that were used to classify the location of contact between a Berghia and its prey. Photo taken by Phoenix Quinlan. B) Plot showing the proportion of appetitive responses that followed a contact between a Berghia and its anemone prey. The error bars represent the standard error. Letters are compact letter display showing the contact locations that were significantly different from one another. A dashed black line represents 0.5. C) Plot showing the proportion of appetitive responses aggregated across all contact types for each FDL. Error bars represent the standard error. D) Plot representing the total number of oral tentacle taps (OT taps) per trial across FDLs. The yellow line represents the fixed effect from the fitted mixed effects model. The lighter yellow shading around the line represents the standard error. The fitted model used a Poisson family. Animal identity nested by the experiment year are included in the model as random intercepts. Diamonds represent the mean. E) A mixed effects model was fit using a Poisson family. The fixed effects were FDL and the state following the approach bout (avoid or feed). Animal identity nested by the experiment year are included in the model as random intercepts. The yellow line represents the fixed effect from the fitted mixed effects model. The lighter yellow shading around the line represents the standard error. Diamonds represent the mean. i. Boxplots representing the average number of OT taps per approach bout that occur during an approach bout that results in feeding. ii. Boxplots representing the average number of OT taps per approach bout that leads to the animal avoiding their prey.

To further dissect how food deprivation influenced exploratory touches, we looked more closely at the number of OT-taps that preceded either feeding or avoiding. The average number of touches per approach bout resulting in feeding was higher than the number of touches before avoiding (b = 1.41, SE = 0.12, z = 12.06, p < 2e-16). Additionally, there was a significant interaction effect of the length of food deprivation on the number of OT-taps that preceded feeding or avoiding (Figure 5 Ei,ii; GLMM; B = −0.16, SE = 0.042, z = −3.80, p = 0.00015). The number of OT-taps during approach bouts that led to feeding was higher and not influenced by hunger, however the number of OT-taps before avoiding increased with food deprivation indicating that the threshold for avoidance is higher in more food-deprived animals.

### Evaluation of contact cues may depend on the stimuli present

To tease apart the chemotactile cues from their prey, gelatin-coated probes were fabricated containing 100% gelatin (tactile control), 10% v/v ground lyophilized anemone with most acontia and nematocysts removed (Flesh probe) and 50% v/v nematocyst and acontia laden ASW (Toxic probe). These probes were applied to the faces and oral tentacles of slugs fixed in place and their behavioral responses were recorded. Contact with the proboscis showed a positive trend (β = 0.211, SE = 0.133, z = 1.59, p = 0.113; Supplemental Table 3), this was not statistically significant. Similarly, contact at the proximal oral tentacle did not differ significantly from distal tentacle contact (β = −0.124, SE = 0.100, z = −1.23, p = 0.219; Supplemental Table 3). This suggests that although contact with the proboscis may have a higher likelihood of triggering feeding behavior, the effect is not strong when accounting for probe type and FDL. Additionally, the proportion of appetitive response by contact location differed from spontaneous contacts and were close to 0.5 regardless of location on the face and oral tentacles of contact (Supplemental Figure 6) This likely indicates the process of pinning them down and or the method of contact dampened appetitive reactions form the slugs.

Probe type significantly influenced the proportion of appetitive responses, while FDL and its interaction with probe type had more nuanced effects (Supplemental Figure 6). Compared to sated animals, the toxic probe under 7-day deprivation elicited significantly more appetitive responses (β = 0.540, SE = 0.161, z = 3.35, p = 0.0008; Supplemental Figure 6). Similarly, the flesh probe under 7-day deprivation also produced a significantly higher response rate (β = 0.352, SE = 0.160, z = 2.20, p = 0.028; Supplemental Table 3). The flesh probe under 0-day deprivation showed a marginal increase (β = 0.269, SE = 0.144, z = 1.87, p = 0.062; Supplemental Figure 6). No significant effects were detected for other combinations of probe type and deprivation. These findings indicate that the response to anemone infused gelatin probes toxic, particularly following food deprivation, are more likely to elicit appetitive behavior than pure gelatin probes.

## Discussion

The nudibranch *Berghia* feeds exclusively on prey that injures them. Here, we confirmed that *Berghia* receive nematocysts in their skin when hunting the sea anemone *Exaiptasia*. We then examined the effect that food deprivation length has on how *Berghia* approaches its prey. All *Berghia*, regardless of FDL, approached their prey. Most of these approaches resulted in contact, which caused injury and risk of capture. Hungry *Berghia* spent more time feeding and orienting towards their prey. Sated *Berghia* generally did not feed. Thus, the contact with prey after initial attraction is the decision point for whether to feed or avoid. The threshold for avoidance was adjusted with hunger such that it took fewer exploratory touches, presumably perceived as aversive, to cause the slug to give up and turn away from the anemone. The results argue against a simple sensory gating model, where peripheral inputs are dampened, and instead support models of central gain control or motivational filtering. Unlike leeches, where feeding suppresses tactile responses (Gaudry and Kristan, 2009; Womelsdorf *et al*., 2014), *Berghia* continues to respond to tactile inputs while feeding and integrates them differently depending on hunger state.

Nudibranchs tend to specialize on particular clades of cnidarians, with some specializing on a single species (McDonald and Nybakken, 1997; Todd, Walter and Daviee, 2001; Goodheart *et al*., 2017). Although all nudibranchs are carnivorous, some species are more aptly compared to host plant-insect interactions due to the behavior of their prey, rather than predator-prey interactions (Cronin *et al*., 1995). However, the *Berghia*–*Exaiptasia* interaction represents a true predator-prey dynamic. *Exaiptasia* tentacles are mobile and it actively defends itself using tentacle movements and acontia ejection, making prey encounters dynamic and risky (Lam *et al*., 2017). *Berghia* specializes on *Exaiptasia* during all life stages (Carroll and Kempf, 1990). Most Aeolid nudibranchs, like *Berghia*, sequester nematocysts from their prey into their cerata, using them for their own defense (Todd *et al*., 2001; Goodheart and Bely, 2017; Goodheart, Barone and Lyons, 2022). This sequestration may be most effective with consistent, specific prey types. Specialization may thus aid resource partitioning, facilitate efficient digestion and detoxification, and enable the development of precise behaviors for prey detection and handling (Bloom, 1981; Edmunds, 1983).

While other studies assume that nematocysts are causing injury and damage to their predators, we were able to directly quantify and visualize nematocysts embedded in a predator following contact (Figure 1). This demonstrates that nematocyst discharge results in physical contact and injury, providing direct evidence of predation costs. The number of contacts is positively correlated with the number of nematocysts observed (Figure 1D), indicating that most interactions with the anemone lead to nematocyst firing. Furthermore, we were able to directly link contacts that resulted in strong aversive responses (e.g., 180-Turn) to dense nematocyst discharge, for example, a contact on the tail that caused the slug to stop feeding and the presence of five nematocysts embedded in the tissue (Figure 1E,F,G). This indicates that the mucous of *Berghia* is not inhibiting nematocyst firing on the oral tentacles despite their specialization, unlike what has been observed in other anemone-specializing nudibranchs (Greenwood *et al*., 2004). Combined with the behaviors following contact—such as pulling away briefly (oral tentacle tap) or sustained withdrawal—this suggests that the slugs are perceiving and responding to nematocysts from their prey.

*Berghia* may have evolved immunity to specific biochemical components of the anemone’s nematocysts. However, it must still detect prey-related cues and remain sensitive to the mechanical damage these stinging cells inflict. Anemone venoms are comprised of many different component peptides with diverse functions (Delgado *et al*., 2022; Fu *et al*., 2022; Michálek, King and Pekár, 2024), some of which may mimic essential neuropeptides that *Berghia* must still respond to—for example, insulin-like (Safavi-Hemami *et al*., 2015, 2016; Ashwood *et al*., 2022).

Although our study was confined to controlled laboratory conditions, naturalistic environments likely increase the risk of injury or predation due to water flow and greater exposure, which could detach slugs from the substrate and facilitate capture. That said, it can be hard to estimate the rate of injury in natural populations, though some studies have done so (Subasi *et al*., 2024). This species has not been studied in natural populations, so our knowledge of their distribution and behaviors in the wild remains limited.

Previous work from our lab found that *Berghia* choose to feed in groups (Otter, Gamidova and Katz, 2025). However, collections of *Berghia* in the wild are often limited to one or two individuals found across a large area (J. A. Goodheart, personal communication), suggesting that group feeding may not be the only strategy employed in nature. This behavioral flexibility may reflect an ecological adaptation to environments where prey are patchily distributed and conspecific encounters are rare. Together, the evolution of prey specialization, tolerance to nematocyst discharge, and context-dependent feeding behavior illustrate how *Berghia*’s ecology and evolution are tightly shaped by the risks and constraints of its interaction with a single, defensive prey species.

### Distance cues cause all animals to approach and evaluate their prey

The presence of prey alone was sufficient to cause *Berghia* to orient and move towards their prey, regardless of hunger state (Fig. 3D, 5C, E). Although other cues may contribute to this behavior, nudibranchs are thought to rely primarily on olfaction to hunt. Studies from our lab have confirmed that rhinophores (the distance chemoreception organ) are required for finding food in a Y-maze (Quinlan, 2024). The behavior of *Berghia*, like some other predatory nudipleurans, is clearly directed motion following detection of prey at a distance (Wyeth and Willows, 2006).

Long range cues from *Exaiptasia* were inherently attractive to *Berghia,* regardless of hunger state. When compared to an empty arena, the presence of prey led animals to orient more frequently and directly toward it, as measured through average heading angle (Figure 3, Supplemental Figure 4). Across all FDLs, animals consistently moved toward their prey (Fig. 3D, 5C, E). Although approaching a sea anemone poses a risk of injury, this baseline attraction may serve functions beyond foraging. For instance, it could promote aggregation to increase reproductive opportunities, enable cooperative or social predation, or guide animals toward locations suitable for egg laying (e.g., near food or shelter). Additionally, it has been hypothesized that innate preferences for visual landmarks are a useful substrate to overlay learning of additional navigational routes (Goulard *et al*., 2021; Buehlmann and Graham, 2022). Olfactory “landmarks” could also play a role in navigation and the odor gradient from prey may also allow slugs to remain in close proximity with a sense of direction and distance (Steck, Hansson and Knaden, 2009; Fischler-Ruiz *et al*., 2021). Multimodal integration is typical in predatory sequences of many animals, with some switching dominant sensory reliance across phases (Lan *et al*., 2012; Gardiner *et al*., 2014). Although *Berghia*’s reliance on olfaction appears primary, the flexibility of sensory hierarchy in response to hunger remains underexplored. In other systems, sensory hierarchies are plastic, shifting with context, ecology, and internal state (Gardiner *et al*., 2014; Kuball *et al*., 2024).

Although long-range cues drive approach, it is more difficult to explain why *Berghia* typically proceeds to make contact, especially given that contact is associated with nematocyst injury (Fig. 1). Nonetheless, distance cues do not only attract animals to the vicinity of prey, they also lead to direct tactile evaluation through chemotactile exploration using the oral tentacles (Fig. 5D–F). Regardless of hunger, animals approach prey and initiate evaluation through contact.

They do so repeatedly across a trial, engaging in multiple approach bouts (Fig. 5A). However, hungry animals differ in that they spend more time per approach bout and return more quickly following avoidance (Fig. 5B–D). The latency between contact and feeding decreases with food deprivation (Fig. 5F), suggesting that hungry animals are quicker to commit to feeding after evaluating prey. The decision to feed or avoid must therefore occur during the contact phase. The decision to feed likely reflects internal processing of contact-related sensory input. These findings are consistent with a broader pattern in specialist feeders, such as *Drosophila sechellia*, where peripheral and central mechanisms shift to enhance responsiveness to preferred stimuli (Chen *et al*., 2024). In *Berghia*, this may reflect a similar adaptation allowing efficient evaluation of familiar but risky prey.

### Hunger shifts the threshold for avoidance in response to contact cues

Once contact is made, the behavioral response of *Berghia* depends strongly on the location of the contact. The probability of an appetitive response is determined by whether contact occurs on the oral tentacles and proboscis, and this pattern is consistent across hunger states (Fig. 6A,B). Contacts elsewhere on the body (head, lips, or body) are more likely to result in avoidance or no response, reflecting responses anemone-initiated contact rather than self-directed exploration. Additionally, when we specifically tried to disentangle cues from their prey by dissociating touch cues from chemical cues of the toxins, nematocysts and secretions, we found that hunger was not sufficient to increase the probability of an appetitive response above 50% for contact with the proboscis, distal oral tentacle or proximal oral tentacle (Supplemental Figure 6). This contrasts with the findings from spontaneous contacts where those parts of the body were likely to have an appetitive response (Figure 6A) regardless of food deprivation. It is possible that the process of fixing them in place dampened appetitive responses.

This distinction between contacts between the anemone and motile sensory appendages such as the oral tentacles and proboscis likely indicates that there is a difference between active sensation/touch and passive contact. Active touch sensing refers to sensation derived from the active movement of the sensory appendage rather than passive contact between the stimulus and the sensor. Often motile sensors are associated with active sensing that can provide additional information quickly as a behavior is occurring (Prescott, Diamond and Wing, 2011). When the anemone’s movement results in contact between the slug and the anemone, this results in aversive reactions to contact. The fact that hunger does not alter this spatial pattern suggests that hunger modulates decision thresholds downstream of sensory localization, perhaps by shifting how information from contact is integrated rather than altering its encoding. This supports the idea that hunger does not suppress peripheral sensory responses, but rather changes how those signals are interpreted centrally. This aligns with neuromodulatory models of state-dependent decision making where hunger alters thresholds without inhibiting detection (Crossley, Staras and Kemenes, 2018).

When specifically examining the main exploratory behavior (OT-taps), we found that the mean number of OT-taps per approach bout increases with food deprivation (Fig. 6C), indicating more active sensing and evaluation of the prey. Interestingly, however, hunger does not affect the number of OT-taps in bouts that precede feeding (Fig. 6D). Instead, the difference emerges only in bouts that precede avoidance: sated animals perform fewer OT-taps before deciding not to feed (Fig. 6E). These findings suggest that hunger does not increase the amount of sensory input required to commit to feeding (i.e., more OT-taps). Instead, they are more willing to persist through aversive feedback before deciding to avoid. During each OT-tap, animals make direct contact with their prey and are likely stung by nematocysts (Figure 1). Yet, hungry animals perform more of these taps before aborting the approach. This suggests that hunger shifts the threshold for avoidance. In this context, hunger may alter how aversive stimuli are weighted relative to appetitive drive, enabling animals to persist in evaluation or feeding behaviors that would otherwise be terminated due to risk or injury.

As a specialist predator, studies of predatory decision making in *Berghia* are different from those in other non-specialist systems. Many decision-making paradigms specifically investigate uncertainty, ambiguity and recognition (Inagaki, Panse and Anderson, 2014; Filosa *et al*., 2016; Rengarajan *et al*., 2019), in this case *Berghia* can sense their prey and through the active evaluation, they are likely not reducing uncertainty about what they are eating, but rather evaluating injury and risk. This is consistent with models where hunger lowers the cost threshold rather than enhancing stimulus quality. That is, sated animals avoid more quickly not because they feel the sting more acutely, but because the aversive weight of the same stimulus is interpreted differently. Overall, the predatory sequence in *Berghia* seems to involve distance detection through olfaction, local evaluation via contact (OT-taps), and decision making based on a balance of aversive and appetitive cues.

*Berghia*, like many other animals, displays a repertoire of stereotyped movements. Many of these movements can be aligned with ethograms described for other predatory nudipleurans (Table 1, Supplemental Table 1). The feeding and foraging behaviors of herbivorous gastropods tends to be similar across species, while carnivorous gastropods show more variation (Elliott and Susswein, 2002). That said, most available nudipleuran ethograms are for generalist predators or herbivores and lack strong empirical validation of those considered aversive or appetitive. For example, oral tentacle withdrawal is described as an aversive response to anemones in the closely related Aeolid, *Aeolidia papillosa*, however we found that this behavior, contrary to what was expected, was most likely appetitive (Edmunds *et al*., 1976). This could be a species difference, or a result of the careful observation and sample size facilitated by laboratory culture of Berghia.

We used multiple human-raters to analyze discrete behavioral responses and contacts with the anemone, however this method is time intensive and susceptible to bias even with blind scoring, especially for subtle behaviors (Marsh and Hanlon, 2007; Tuyttens *et al*., 2014). The advancement of tools for automated machine-learning based tracking of key points, such as Deeplabcut, has made it relatively straightforward to track most animals (Pichler and Hartig, 2023). Soft-bodied invertebrates pose additional challenges for accurate key point detection that can be overcome through sufficient training and carefully standardizing the viewing angle of the animal (Sivitilli *et al*., 2023). Automated, markerless pose estimation in soft-bodied animals like *Berghia* is challenging due to their flexibility and the presence of numerous appendages such as cerata and oral tentacles. These features cause many occlusions of key body parts that require extensive training data to overcome. Since the analysis of this study, advanced segementation models such as SAM2 have begun to be used in biological research and should be used in future studies of this and similar animals (Mele *et al*., 2026). In this study we employed a strategy that combined automated tracking of limited key points informed by our understanding of predatory behavior in this species with thresholds derived from human observations. Our method demonstrates the feasibility of automated behavior quantification in soft-bodied invertebrates with variable appendages and emphasizes the importance of leveraging human intuition about animal behavior to determine useful measures to quantify. We were able to conduct direct, repeatable measurements of behavior using this approach.

### Potential neural mechanisms involved in this satiety-dependent decision making

Hunger powerfully shapes behavior and perception of animals and can act at peripheral and central levels to do this. Satiety signals are often widespread and impact multiple neural substrates (Tierney, 2020; Flavell *et al*., 2022). In *Drosophila*, for instance, state-dependent modulation occurs both in the antennal lobe and in integrative regions like the mushroom body (Tsao *et al*., 2018; Lin, Senapati and Tsao, 2019). In nudipleurans such as *Pleurobranchaea*, modulation occurs centrally and peripherally (Hirayama and Gillette, 2012; Brown *et al*., 2018). Mechanistically, this can involve neuromodulators like serotonin or NPY, which reconfigure network dynamics (Jing *et al*., 2007; Marder, O’Leary and Shruti, 2014).

In *Berghia*, the persistence of tactile reflexes despite hunger suggests central reweighting rather than presynaptic inhibition. It is possible that these stable responses across hunger states are a result of reflexes, which could be true for behaviors like bristling (Brown *et al*., 2024), however while the proportion of appetitive responses was stable, the specific behavior that occurred was not stable, just the class of behavior. Additionally, acute responses to pain are often separate from integrated responses such as turning away and leaving which can be hunger dependent (Alhadeff *et al*., 2018). This shift in cost tolerance with hunger could reflect neuromodulatory changes that alter the salience of aversive input or suppress avoidance circuitry. Hunger state may modulate sensory processing or decision-making thresholds via mechanisms such as serotonergic or dopaminergic gain control, enabling animals to persist in behavior despite negative consequences. Such central reconfiguration has been observed in other mollusks (Hirayama and Gillette, 2012; Crossley, Staras and Kemenes, 2018), where feeding-related interneurons shift network dynamics. It is likely that similar processes occur in *Berghia* to shape approach avoidance decision making where central mechanisms such as motivational filtering or gain control are used.

## Methods

### Animal Care

A culture of *B. stephanieae*, originally sourced from Salty Underground (Crestwood, MO, USA) and Reeftown (Boynton Beach, FL, USA), was maintained in the lab as described elsewhere (Otter, Gamidova and Katz, 2025). *B. stephanieae* were housed communally in groups of 5–15 individuals in 1-gallon acrylic aquariums filled with artificial seawater (ASW; Instant Ocean, Blacksburg, VA, USA) with a specific gravity of 1.020–1.022 and pH of 8.0–8.5, under a 12:12 light-dark cycle at 22–26°C. *Exaiptasia diaphana* (Carolina Biological Supply Co., Burlington, NC, USA) were housed in glass aquariums with ASW under the same conditions as *B. stephanieae*, with additional filtration. *E. diaphana* were fed twice weekly with freshly hatched *Artemia nauplii*. The nauplii were hatched every two days by placing 2.5 g of freeze-dried *A. nauplii* eggs into an aeration chamber with fresh ASW. *B. stephanieae* were fed twice weekly by placing two *E. diaphana* individuals into their home tank, unless otherwise noted.

### Experiment 1: Quantifying injury

#### Behavior Experiment

To test the hypothesis that anemones are dangerous because they fire nematocysts into their predators, we used confocal microscopy to directly visualize the skin of *Berghia* following interactions with *Exaiptasia.* Adult slugs were selected for this experiment, with smaller individuals prioritized to facilitate imaging. The experimental setup included a self-leveling mat to minimize vibrations and was backlit using a white LED lightboard for even illumination. Prior to testing, slugs were acclimated for five minutes in an arena identical to the testing arena placed on a white LED lightboard. Simultaneously, the anemone was acclimated in the testing arena under the same lighting conditions. Following acclimation, slugs were introduced to the testing arena and filmed for 15 minutes. Videos were analyzed using Behavioral Observation Research Interactive Software (BORIS v. 8.24; Friard and Gamba, 2016) to quantify slug behavior and count the number of contacts between the slug and the anemone.

#### Fixation and Tissue Preparation

To visualize the skin of the *Berghia,* slugs were anesthetized immediately after removal from the experimental arena in 4.5% MgCl₂ in ASW. The anterior-most part of the face was removed using fine dissection scissors (Fine Science Tools) by making a cut just posterior to the oral palps. The oral tentacles were spread apart using 0.1 mm minutien pins (Fine Science Tools) to minimize damage, secured in an “X” formation without piercing the skin. Faces were fixed in 4% paraformaldehyde (Electron Microscopy Sciences) in 4.5% MgCl₂, which allowed re-imaging as necessary. During fixation the body parts were pinned in Sylgard-lined Petri dishes to ensure the position would facilitate mounting and analysis. If the anemone touched the slug’s tail, the tail was also removed and fixed following the same protocol. After overnight fixation at 4°C, samples were washed with 1X phosphate-buffered saline (Fisher Scientific) and mounted on a glass slide using 100% glycerol.

#### Confocal Microscopy and Image Analysis

Nematocysts embedded in the skin of *Berghia* were visualized using a Zeiss LSM 700 laser-scanning confocal microscope. Imaging was performed using a 40X oil immersion objective with differential interference contrast (DIC) to enhance structural detail. Each animal had its left oral tentacle, right oral tentacle, and lips imaged, with later iterations also including the tail. Z-stacks were acquired at a depth of 100 µm. Images were processed in ImageJ and divided into a 100 µm grid. Within each section, nematocysts were identified based on their characteristic ovate shape and counted using the ImageJ cell counter plugin. The total number of nematocysts per sample was recorded for analysis.

### Experiment 2: Binary food deprivation

To compare the behavior of hungry and sated slugs, we quantified the behavior of slugs after 0- and 7-days of food deprivation. Slugs were housed individually in 355 mL deli cups labelled to maintain identity. After each experiment, slugs were provided with three fresh anemones overnight, and cups were cleaned after 24 hours. No slugs consumed three anemones within 24 hours, so we considered providing three anemones overnight *ad libitum* feeding. Slugs remained in isolation between experiments. Before testing, each slug was fed *ad libitum* for 24 hours, then isolated and food-deprived for seven days unless assigned to the 0-day food deprivation condition, in which case they were provided anemones the day before their trial. Trials were conducted seven days apart. Slugs were divided into four groups: (1) 15 slugs tested at 0 days food-deprived twice to assess repeated testing effects in the absence of food deprivation, (2) 15 slugs tested at 7 days food-deprived twice to examine repeated testing effects under prolonged deprivation, (3) 15 slugs tested at 7 days food-deprived first, followed by 0 days food-deprived, and (4) 15 slugs tested at 0 days food-deprived first, followed by 7 days food-deprived, to compare behavioral differences within the same individuals while controlling for order effects.

Experiments were conducted in a clear acrylic arena (7.62 × 7.62 × 2.54 cm) placed on a black self-leveling ¼” thick EVA foam pad (Grainger) to reduce vibration. The arena was illuminated from below by a white LED lightboard for even illumination. The edges of the arena were covered by white opaque window film to eliminate external visual features. Filming was conducted from above using a Canon EOS 80D camera with a Canon EF 50mm f/1.8 STM lens. Each trial lasted about 15 minutes, with the first 11 minutes analyzed, beginning after the slug righted itself in the arena. The anemone was placed approximately in the center of the arena and allowed to acclimate for five minutes while the slug simultaneously acclimated in an identical separate arena under the same lighting conditions. The slug was then transferred to the test arena using a plastic transfer pipette with the distal 2 cm cut off to widen the opening. Filming proceeded undisturbed for 15 minutes before the slug was removed and returned to its housing cup.

To compare behavior in an arena with prey to the behavior of *Berghia* in an empty arena, the setup was identical except that the arena was placed directly on a black self-leveling pad and surrounded by four white LED lightboards for uniform illumination without backlighting. After a five-minute acclimation, slugs were placed in the arena and filmed for 15 minutes. Each slug was tested twice, once at 0 days food-deprived and once at 7 days food-deprived. Half were tested in the 0-day condition first, and the other half in the reverse order. This experiment allowed us to detect which behavioral changes were due to prey presence as opposed to space utilization difference in hungry and sated slugs.

To compare the behavior of *Berghia* in an empty arena and with prey to an arena with a novel object in the center, the setup remained unchanged except that an opaque white marble was placed at the center of the arena. Ten slugs were tested at 0 days food-deprived, and a separate set of ten slugs was tested at 7 days food-deprived. No slugs were tested multiple times in this experiment. Each trial lasted 15 minutes. This allowed us to be sure the behavior with prey is specific to prey and not just due to the presence of an object.

### Experiment 3: Serial food deprivation

To test individual *Berghia* across multiple FDLs we performed experiments as in the other experiments with a few key differences. The setup matched the empty arena experiment, using four LED lightboards and a black self-leveling pad to minimize vibration. The experiment was conducted in two cohorts: Summer 2021 and Summer 2022. In each cohort, 30 slugs were individually housed in plastic deli cups (500 mL) and split into six groups to counterbalance order effects across FDLs (0–5 days). The second cohort included 33 animals to account for missing data from the first cohort. Each experiment began with a five-minute acclimation, followed by 15 minutes of filming. Only the 13 minutes after the slug righted itself were analyzed. Animals missing usable data for at least four trials were omitted due to illness or death during the experiment or camera failure resulting in fewer than 13 minutes of usable footage. After each trial, slugs were given ad libitum access to anemones for 24 hours. Depending on their assigned group, they were then food-deprived for 0–5 days before being tested again. Each slug was tested six times across five trials.

### Experiment 4: Controlled touch

#### Data Acquisition

To test the prediction that *Berghia* reacts to the toxins from *Exaiptasia* differently than to the other cues present during contact, we disassociated the cues by making artificial probes to touch the *Berghia.* Behavioral responses to chemotactile stimuli were recorded using a Leica M165C microscope equipped with a Leica IC90E integrated CMOS camera, which was wirelessly operated using the Leica EZ4 W & ICC50 W Remote Control and displayed on a Dell 24” monitor. A Siskiyou micromanipulator, controlled by a Siskiyou MC1000e push-button controller and MC1000e controller outlet, was used to deliver precise tactile stimulation. The micromanipulator allowed for longitudinal and oblique movement at a rate of 1.7 mm/s and 300 µm/s, respectively. An AmScope LED 6WD spotlight was positioned to optimize visibility during motion capture.

#### Probe Fabrication

Gelatin-coated probes were designed to present chemotactile stimuli while ensuring sufficient structural integrity for repeated contact with the oral tentacles of *Berghia*. To construct the probes, the tip of a 1 mL SM bulb pipette was heated with a BIC lighter until malleable, then stretched and twisted to a final diameter of approximately 0.5 mm. The excess was removed, resulting in a 170 mm probe.

Stimuli preparation involved separating *E. diaphana* tissue from nematocysts and acontia. Anemones were vortexed in 2 mL of artificial seawater (ASW) within 5 mL conical-bottom tubes for 60 seconds, a method used by *Exaiptasia* venom researchers to extract venom (J. Macrander, personal communication). Vortexing caused the anemones to fire nematocysts, releasing toxins into the water and eject acontia which could then be separated from the body of the anemone. The anemones were then removed with forceps and transferred to a separate tube, while the nematocyst, toxin and acontia-laden water was collected separately. The separated anemones were lyophilized using a VirTis Genesis Pilot Freeze Dryer (Scientific Products, Warminster, PA, USA) and ground into a powder, which was mixed with a thickened gelatin solution (10% w/v with 50% ASW and 50% hot distilled water) to create an anemone-infused gelatin (10% w/v lyophilized anemone). Meanwhile, the nematocyst, toxin and acontia-containing ASW was used to make gelatin (50% toxin and nematocyst-laden ASW and 50% hot ASW) to create a toxin-infused gelatin solution (50% v/v toxin and nematocyst-laden ASW). The mixtures were visually inspected for nematocysts to ensure the toxin-containing liquid had them and the anemone powder did not. Probes were dipped into the respective gelatin mixtures and allowed to solidify before use, ensuring consistent application of either the anemone-derived powder or acontia-infused gelatin for experimental trials.

The toxin and nematocyst laden water toxicity was confirmed using an artemia toxicity assay (Chan *et al*., 2021). Briefly, *A. nauplii* were hatched as described in the animal care section. Four solutions were prepared using ASW with 0%, 5%, 25% and 50% v/v toxin and nematocyst laden water mixed with ASW. Four replicates of 10 animals were used to determine the toxicity of each concentration after 1 and 24 hours. All solutions containing the toxin and nematocyst laden water had 100% lethality after 24 hours and the 50% solution was lethal within one hour (Supplemental Figure 7). The lyophilized anemone could not be used for this assay because when in contact with water it becomes gelatinous and artemia could not survive in a solid.

#### Behavioral Assay

Slugs were secured for testing using two Minutien pins (Fine Science Tools) placed between the rhinophores and the heart and in the posterior body. A 4-axis micromanipulator (SD Instruments) was used to deliver gentle tactile stimulation to the distal oral tentacle, proximal oral tentacle, and lips. Each body part was stimulated approximately five times per trial using the gelatin-coated probes.

Behavioral responses and probe contacts were manually scored using BORIS. Each video was analyzed in five passes: (1) probe position (medial or lateral relative to the oral tentacles), (2) number of contacts (including location of the contact: distal/middle/proximal oral tentacles, proboscis, lips and head), (3) protruding behavior (Table 1), (4) slug behaviors (Supplemental Table 4), and (5) an independent blind review of behaviors and contacts. Raters were blinded to both the probe type and the food-deprivation condition of the slug. Slugs used in this experiment were either 0- or 7-days food deprived, consistent with prior experiments.

### Video analysis & behavior quantification

Each video was processed using a custom Python script utilizing *ffmpeg* to standardize the format and reduce file size. If necessary, videos were converted to MP4 format, and audio tracks were removed. Videos were then re-encoded using the H.265 codec (*libx265*) with a constant rate factor (CRF) of 18, balancing file compression and quality. The “veryfast” preset was used to optimize processing speed while maintaining efficient compression. A single frame from each video, where the slug’s body and oral tentacles were fully extended, was used to measure body length, oral tentacle length, and arena dimensions. If applicable, marble diameter and center coordinates were also recorded. ImageJ was used to annotate these features, including capturing pixel-space coordinates for the arena’s corners.

To estimate spatial relationships, the Euclidean distance between arena corners was measured, and the average side length of the 76.2 mm square arena was used to calculate the pixel-to-millimeter ratio. This ratio was applied to convert slug length, oral tentacle length, and anemone radius to millimeters. The anemone radius was estimated from its measured area. Since anemones were not fixed in position, the maximum possible distance between the slug and the anemone was calculated by determining the Euclidean distance between the anemone’s mouth and each arena corner, selecting the largest value.

#### Manual Event Logging

Videos from Experiments 1, 3, and 4, as well as videos containing prey in Experiment 2, were manually scored using BORIS software. The ethograms used for Experiments 1, 2, and 3 are available in Supplementary Table 3, while a streamlined ethogram was used for Experiment 4 (Supplementary Table 4).

Raters were trained using five sample videos, during which at least two other raters reviewed the videos together to ensure agreement. Following training, all raters independently scored the same set of 10 videos to assess inter-rater reliability. Each video was reviewed twice: once to encode contacts between the slug and anemone, and a second time to document discrete behaviors of both organisms. The exported CSV files were processed using custom R scripts.

#### DeepLabCut Tracking

To track the movement of slugs and anemones, DeepLabCut (Mathis et al., 2018) neural networks were trained separately for different experimental conditions. One network was trained to track slugs and anemones in Experiments 1 and 3, another for slugs and anemones in Experiment 2, and a third for slugs in arenas without anemones in Experiment 2. Additionally, a separate network was trained to track detailed body points on the slugs. All networks used a ResNet-50 backbone, and tracking outputs were filtered using a spline filter with a window length of 5 to reduce noise.

**Figure 6.**
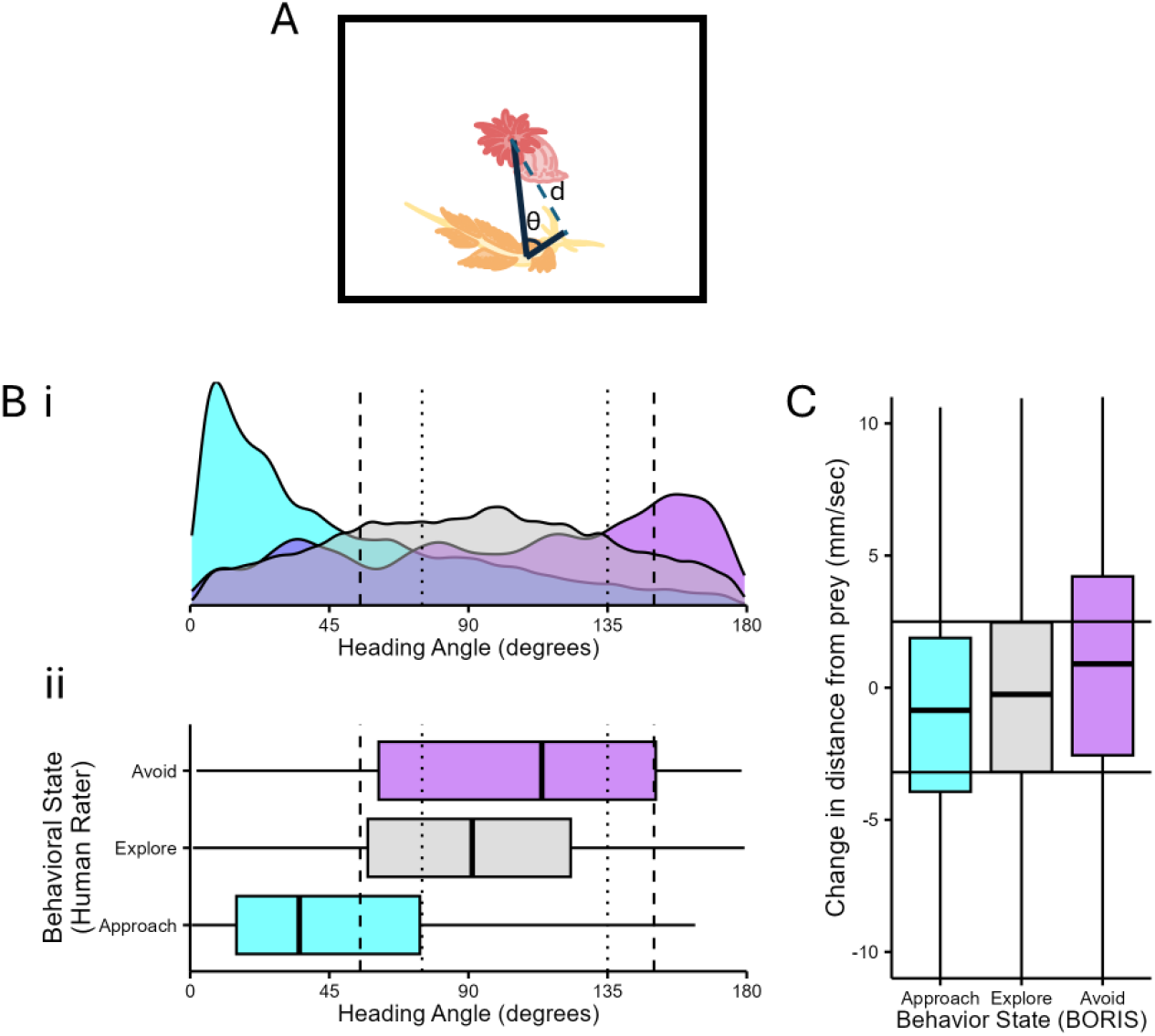
Approach and avoidance behavioral states were defined by heading angle and change in distance from prey. A) A schematic depicting how distance from prey (d) and heading angle (θ) were calculated, for slugs with prey in the arena. We used the heading angles from the frames assigned to each behavioral state by a human rater to determine thresholds for approach and avoidance. The heading angle is defined as the angle formed between the opening of the gastrovascular cavity (henceforth referred to as the “mouth”), the middle of the body of the slug (a white opaque patch located above their heart) and the lips of the slug.B) i. A smoothed density plot of heading angles. ii. A boxplot showing the angles from these same frames. The colors represent Approach (cyan), Explore (gray) and Avoid (purple). The dashed lines represent hard thresholds where any frames with a heading angle above the 3rd quartile of heading angles classified as avoidance (#) or below the 1st quartile of heading angles labeled by a human as exploring are classified as avoid or approach, respectively. The dotted lines represent soft thresholds where heading angles between the hard and soft thresholds for approach and avoid rely on the change in distance from prey to determine approach or avoidance. See the Methods for details. C) The change in distance between the mouth of the slug and the mouth of the prey for each frame classified by a human rater. The 1st and 3rd quartile of frames labeled approach and avoid respectively were used as thresholds for determining if frames with heading angles in the soft threshold ranges were approach or avoidance.

Each slug was tracked at four primary points: lips, head, mid-body, and caudal body. An additional network was trained to track 16 specific points: the lips, head, mid-body, three additional points along the caudal body, tail, and six locations along each oral tentacle (base, proximal, middle, distal, and tip). While this model performed well in some cases, it struggled with occlusions during interactions, limiting its usefulness to short periods without occlusions. A total of 1,580 labeled frames from 107 videos were used to train this model. For the primary networks, training proceeded iteratively through refinement rounds until the train and test errors converged below 5 pixels. After initial training, if more than 20% of frames in a video contained keypoints with a likelihood below 0.8 for the anemone’s mouth, the slug’s lips, or the middle of the slug’s body, additional frames were manually labeled for refinement.

For Experiment 2 with anemones, the network was trained using the resnet_v1_50 backbone and refined over five rounds. The first three rounds used a training fraction of 0.95, while the last two used 0.90. A total of 1,107 labeled frames were used, and the network was trained for 2,980,000 iterations. The network used for Experiments 1 and 3 was refined over 11 rounds with a training fraction of 0.90. Data augmentation included image rotation up to 80 degrees (*imaug* augmentation). This model was trained using 3,361 labeled frames from 215 videos, with a total of 4,780,000 iterations. For tracking slugs in arenas without anemones (Experiment 2), a separate network was trained over three rounds of refinement using 819 labeled frames from 38 videos. This model was trained for 2,500,000 iterations.

Post-processing using custom R scripts (github) to calculate the heading angle, distance from the anemone and change in distance from the anemone. These measurements were used to determine the behavioral state of the slugs as described previously.

#### Automated Behavior State Classification

To quantify the amount of time animals spent approaching or avoiding their prey, we estimated angle and change in distance thresholds based on observations of animal behavior. Human raters utilized Behavioral Observation Research Interactive Software (BORIS; Friard and Gamba, 2016) to annotate the videos, categorizing behaviors as described in Table 1. Throughout the trials, raters were instructed to assign each frame to one of four behavioral states: approach, avoid, feed, or explore. These behavioral states were defined using heading angles that reflected the arena’s geometry. However, the human raters did not measure the heading angles directly but used them as guidelines when annotating behaviors.

The behavioral state assignments made by the human raters were then used to determine the thresholds for automated tracking with DeepLabCut (Nath *et al*., 2019). Specifically, the first quartile of heading angles from frames labeled “Explore” served as a cutoff for the “Approach” state (Figure 1A). Any frame where the animal’s heading angle was smaller than this threshold was considered as “approaching.” Similarly, the third quartile of heading angles from frames labeled “Explore” was used as a cutoff for the “Avoid” state, with any frame where the animal’s heading angle was larger than this threshold considered as “avoiding” (Figure 1 A).

In addition to these hard thresholds, soft thresholds combining both heading angles and the change in distance from the prey were defined. For the “Approach” state, a frame was considered “approaching” if the angle was smaller than the third quartile of heading angles for frames labeled “Approach,” and the change in distance from the prey was more negative than the third quartile of the change in distance from prey for frames labeled “Explore.” Similarly, for the “Avoid” state, a soft threshold was applied that combined the heading angle and change in distance (Figure 1 B). Thus, the criteria for approach were as follows:

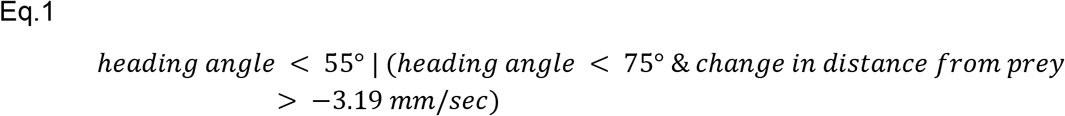

and for avoidance:

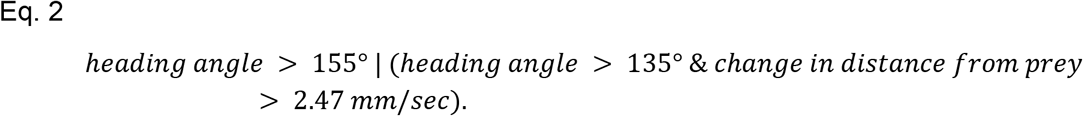

This method enabled unbiased detection of approach and avoidance with clearly operationalized definitions.

To determine the duration of time in each state, it was determined whether the *Berghia* in each frame of every video was approaching, avoiding, or neither. Brief changes in heading unrelated to sustained movement were accounted for by ensuring behavioral states persisted for at least 14 seconds before being classified as a transition. This time was determined by measuring the dwell time and transition frequency for different time intervals and using the geometric mean of the inflection points detected. To validate that this method of direct quantification was comparable to human ratings, the duration of time spent in each behavioral state was compared. Notably, the results were more consistent in food-deprived animals. The primary difference was that human raters were less likely to classify an animal as avoiding, regardless of FDL, and instead categorized them as exploring (Supplemental Figure 2).

The impact of food deprivation on time spent in each behavioral state was compared between the two quantification methods. The statistical relationships were consistent across both methods (Supplemental Table 2), with the exception of time spent avoiding, which the threshold method only detected a decrease in food-deprived animals (Supplemental Figure 3). Since human raters were less likely to classify frames as avoiding, the direct quantification method was more sensitive to this change (Supplemental Figure 2).

The time a slug spent making contact with and interacting with their prey was called “handling time.” Handling time was automatically classified for any frames where the slug was not eating, the heading angles were smaller than 100 degrees and when the distance between the lips and the center of the prey was smaller than the radius of the prey and the length of the oral tentacles.

#### Binary Response Calculation

To determine how different types of contact influenced subsequent behavior, we identified behaviors that followed each contact event within predefined time windows. Contact events were categorized by the specific body part where the anemone and the slug came into contact: body, head, lips, proboscis, proximal oral tentacle, distal oral tentacle. For each contact, we recorded whether a response occurred and, if so, whether it was appetitive or aversive. Behaviors were only included if they occurred within a time window appropriate to their typical latency (short: <1s, medium: <2s, long: <5s), accounting for the slugs’ slow movement (e.g., a 180-Turn may occur with a longer delay). Time windows began 100 ms (about 3 frames) before a contact and ended 100 ms before the next, ensuring each behavior was attributed to a single contact. Behaviors not fitting any window were excluded. When multiple behaviors followed a contact, we calculated the average response and rounded to the nearest 0 or 1 (with 0.5 rounded down) to determine the net response.

### Statistical analysis

All statistical analyses were conducted in R (R Core Team, 2024) using custom scripts available upon request. Specifically, we utilized the following packages for data manipulation and analysis: dplyr (Wickham, François, *et al*., 2023), tidyr (Wickham, Vaughan, *et al*., 2024), data.table (Barrett *et al*., 2024), lme4 (Bates *et al*., 2025), GLMMTMB (Brooks *et al*., 2024), emmeans (Lenth *et al*., 2025), multcomp (Hothorn *et al*., 2025), multcompView (Graves and Dorai-Raj, 2024) and trajr (McLean, 2023). For data visualization, we used: ggplot2 (Wickham, Chang, *et al*., 2024), cowplot (Wilke, 2024), ggpubr (Kassambara, 2023), scales (Wickham, Pedersen, *et al*., 2023), and RcolorBrewer (Neuwirth, 2022).

To examine the relationship between the number of contacts and the number of nematocysts embedded in faces, a linear regression was performed. All statistical tests were two-tailed, with significance set at p < 0.05. Paired and unpaired t-tests were applied where appropriate to compare mean differences, with effect sizes reported using Cohen’s d. For binary food-deprivation experiments, paired t-tests with Bonferroni correction were used to compare within-animal measures across 0 and 7 days of food deprivation. To compare the empty arena, arena with prey, and arena with the marble, we employed Tukey’s Honest Significant Differences test to perform pairwise comparisons of arena types and FDL within those arenas. When the assumptions for a t-test were not met, a Wilcoxon test was used.

For the serial food-deprivation experiments, all data were modeled using generalized linear mixed models (GLMMs) with animal identity and experimental cohort (year) as random effects. The primary fixed effect of interest was FDL, modeled as a continuous variable to capture smooth behavioral changes across days. Model selection assessed additional predictors, including the size difference between anemones and *Berghia*, time of day, and the previous period of food deprivation experienced. Model comparison was performed using the Bayesian Information Criterion (BIC), and the best-fit model always included FDL as the sole fixed effect. For any data that appeared to have a nonlinear trend, we also we compared models that modeled food deprivation as a discrete variable, but it was not a better fit than the model using continuous representation in all cases, indicating that the data is best modeled when food deprivation is treated as a continuous variable. For measures restricted to positive values, a Gamma distribution with a log-link function was used (e.g., distance from prey, average heading angle). Durations were transformed to a (0,1) scale by dividing by the trial length (780 seconds) and adding 0.001 to feeding time before modeling with a beta distribution and logit-link function. This approach was also applied to time avoiding and handling time.

To examine whether food deprivation influenced the likelihood and timing of feeding following contact with prey, we used a mixed-effects Cox proportional hazards model (coxme package in R). This survival analysis approach models time-to-event data while accounting for random effects due to repeated measures. In this case, the time from initial contact to feeding was treated as the response variable, with FDL modeled as a discrete fixed effect (factor), and animal identity nested within experimental cohort (year) included as a random intercept. This structure accounts for non-independence of trials from the same individual and temporal variation across cohorts. Model fit was assessed using the Bayesian Information Criterion (BIC), and hazard ratios were used to interpret the effect of food deprivation on feeding latency. Significance testing was two-tailed with p < 0.05. Full model results, including random effect variances and log-likelihood comparisons, are provided upon request.

## Notes

### Competing Interest Statement

The authors have declared no competing interest.

### Summary of Updates

The previous upload was missing references. Discussion of pose estimation tools was updated.

### Funding

- National Science Foundation: 2227963, GRFP
- National Institutes of Health: U01-NS10863, U01-NS123972, NIH R0Z1NS133654

