## Supplemental Figures and Tables for "Hunger modulates behavioral responses to olfactory and chemotactile cues in the specialist predator of dangerous prey, *Berghia stephanieae*"

Supplemental Information

### Supplemental Table 1

| Supplemental Table 1: Comparison of *Berghia* ethogram to ethograms of related species. | | |
| --- | --- | --- |
| Behaviors | **Description** | **Other Species (description)** |
| Bite | *Berghia* bites; buccal mass can be seen moving outward and then inward | *Aplysia californica:* Biting (Leonard and Lukowiak 1986) |
| Bristle- Lateral | *Berghia*’s cerata extend and move along the lateral axis of the body (coronal/frontal plane in humans) | *Tritonia plebeian:* bristle cerata (did not distinguish subtypes (Allmon and Sebens 1988)  *Coryphella verrucosa:* bristle cerata (did not distinguish subtypes (Allmon and Sebens 1988) |
| Bristle- Medial | *Berghia*’s cerata become erect and move medially towards the center of the body, making the *Berghia* appear thinner in one area or throughout the body. |  |
| Bristle- Rhinal | *Berghia*’s first cerata cluster become erect and move medially and towards the head and rhinophores, often occluding the head and eyes; this is distinct from a head withdrawal because the head remains in place and the cerata move forward to cover it. | *Navanax imermis*: similar to Shrug Parapodia (Leonard and Lukowiak 1984) |
| Curl | *Berghia* laterally flexes its body at the level of the middle of its body until its head is parallel to its posterior body and pauses in this “C” shape; tail and posterior body remain in place | *Aplysia californica:* Balled-up (Leonard and Lukowiak 1986)  *Aplysia californica:* Defensive Withdrawal (Jahan-Parwar and Fredman 1979) |
| Head Wave | With posterior body stationary, *Berghia* bends at the mid body and moves its head laterally and back to midline in both left and right directions | *Aplysia californica:* Head-waving (Leonard and Lukowiak 1986)  *Navanax imermis:* Head-waving (Leonard and Lukowiak 1984) |
| Head Withdrawal | *Berghia* retracts its head towards the middle of its body, cerata may perform a rhinal bristle, eyes and head spot disappear from view, tail and posterior body remain in place | *Aplysia californica:* Head withdrawal (Audesirk and Audesirk 1985; Leonard and Lukowiak 1986)  *Navanax inermis:* Head Withdrawal (Leonard and Lukowiak 1984) |
| Oral Tentacle Tap | Distal end of the oral tentacle makes contact with an object, briefly flexing away (often dorsally), and then immediately returns to a resting position or makes contact again |  |
| Oral Tentacle Withdrawal | *Berghia* retracts one of its oral tentacles flexing dorsally at the proximal end; oral tentacle is pulled behind the lips |  |
| Oral Tentacles “W” Pose | Both of *Berghia*’s oral tentacles flex dorsolaterally at the proximal end such that the tips end up level with the head and for a “W” shape; this pose is held for at least 0.5 seconds |  |
| Oral Tentacles Cross | Both of *Berghia*’s oral tentacles flex dorsomedially at the proximal end such that the tips end up level with the head and in line with the body, often they will form an “X” shape and the tips will cross midline |  |
| Orient | From a neutral heading to their prey or from a directionally opposite heading to their prey *Berghia* turns its head towards their prey and proceeds in that direction with its body following |  |
| Pause | *Berghia* does not move its body in any direction for a period of time; the head still moves slightly and often flexes dorsally off the substate |  |
| Rear | *Berghia* lifts its head dorsally off the substrate for more than 0.5 seconds | *Aplysia californica:* Rearing (Leonard and Lukowiak 1986) |
| Scrunch | *Berghia* retracts its midbody towards the posterior body, while the head and oral tentacles remain out and then relaxes back into a normal state. | *Navanax imermis:* similar to slight head-withdrawal (Leonard and Lukowiak 1984) |
| Start Protrude | *Berghia* everts its buccal mass/proboscis; an “m” shaped structure appears in front of the lips. | *Tritonia diomedea:* When describing “bite -strikes” says lips protruded (Wyeth and Willows 2006) |
| Tail Withdrawal | *Berghia* retracts its tail, such that the tail disappears from view or retracts its posterior body such that its length shortens with movement; only occurs in the posterior body | *Aplysia californica:* Tail-withdrawal (Leonard and Lukowiak 1986)  *Navanax imermis*: Tail-withdrawal (Leonard and Lukowiak 1984) |
| Turn - 180 | *Berghia* lifts its head from the substrate and rotates its head around the middle of its body until it is pointed away from the anemone; returns head to the substrate and moves in the new direction with the posterior body following |  |
| Turn- 360 | *Berghia* turns 360 degrees or more in one direction; entire body completes a circle |  |
| Turn- F | *Berghia* temporarily turns its head 90° away from the prey, while the end of its body does not fully turn along with the head |  |
| Turn- J | From a position facing away from prey, the *Berghia* turns its head 180° towards the prey without its body following after |  |

### Supplemental Figure 1

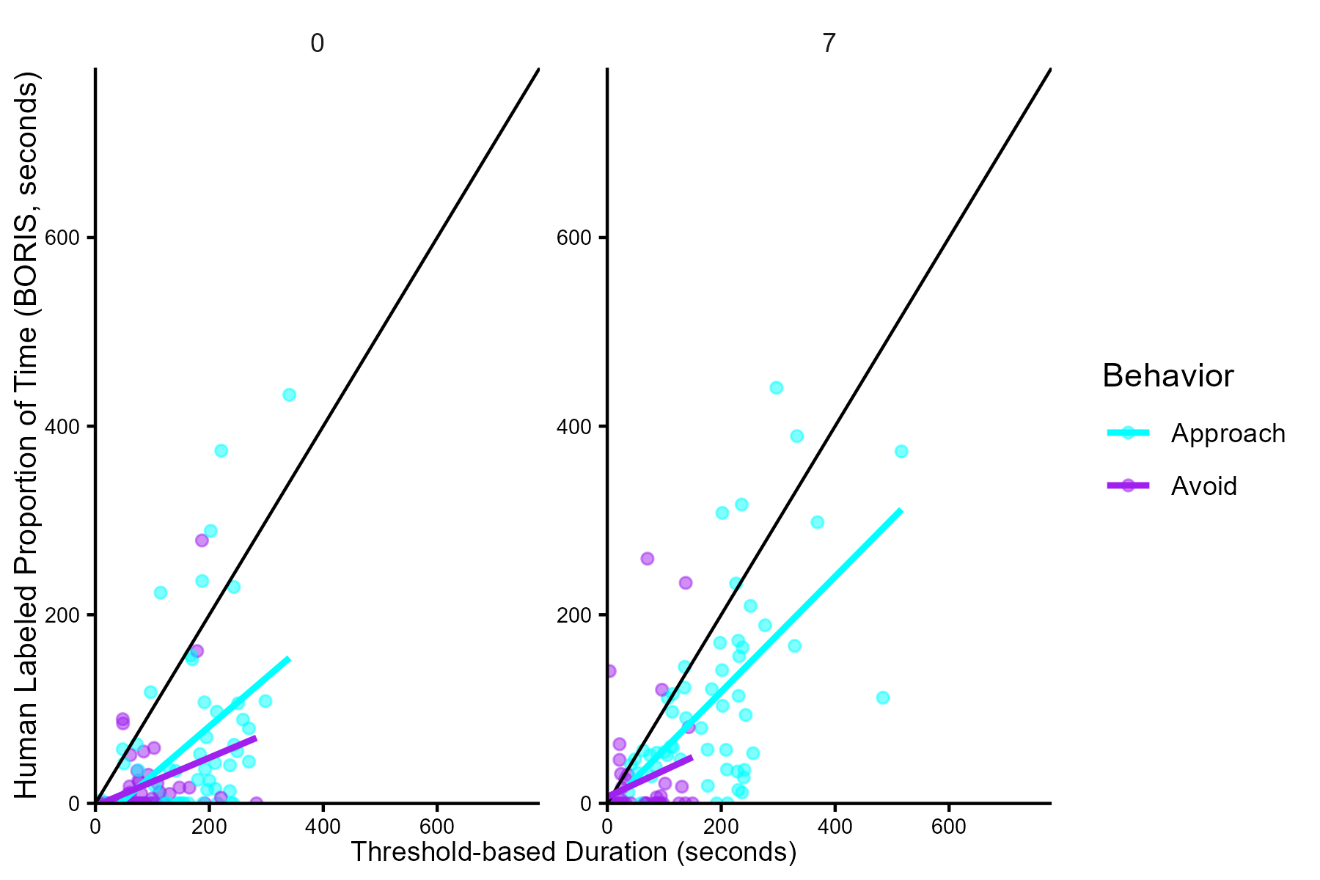
SI Fig 1: Human-raters are less accurate at determining avoidance than approach, especially in sated animals. Plots show the relationship between the duration of time a human-rater assigned to approach (cyan) or avoidance (purple) in sated (left) and 7-day food-deprived animals (right). The black line represents the y=x line which would indicate a perfect match in duration.

### Supplemental Figure 2

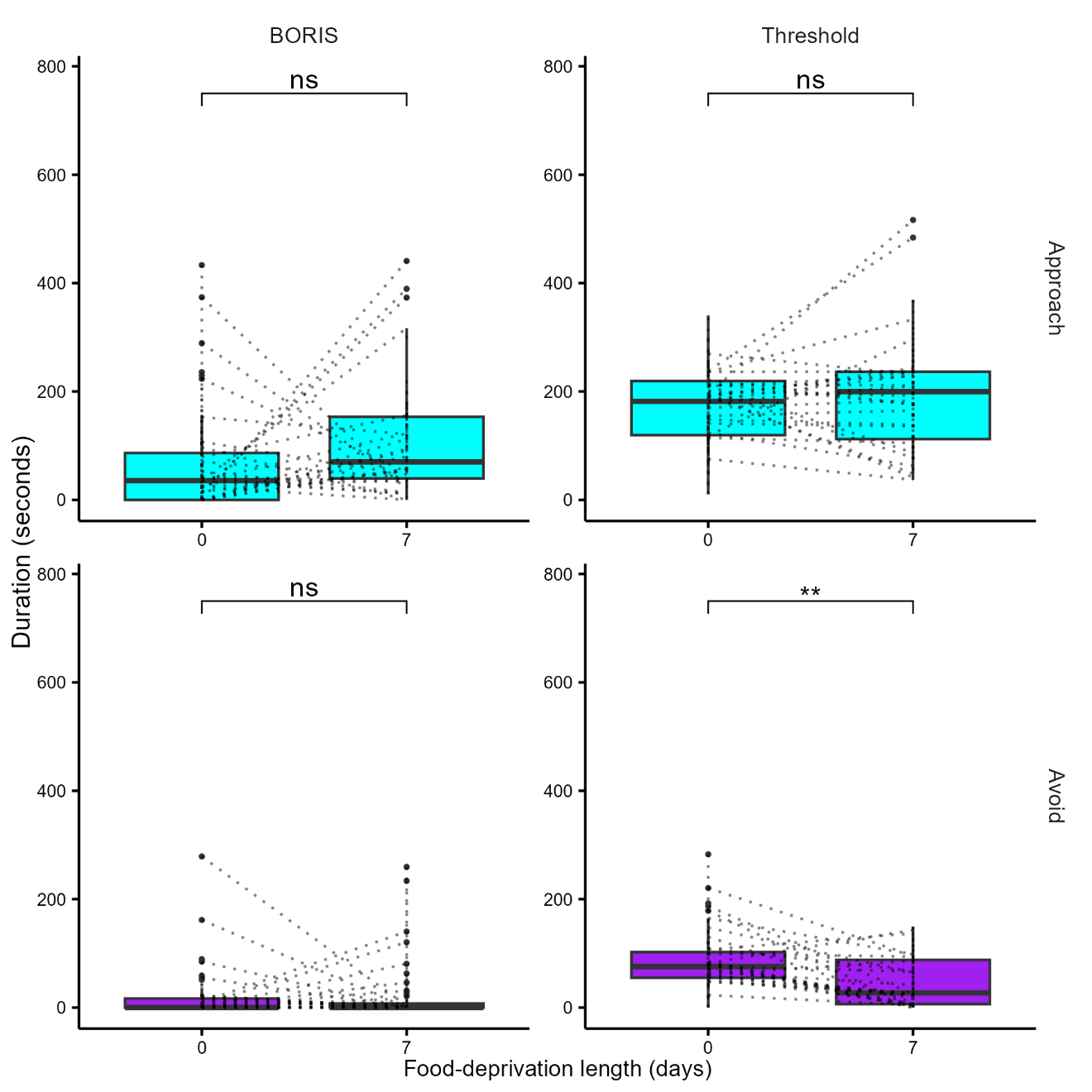

SI Fig 2: The effects of food-deprivation on approach match between analysis methods, however a decrease in duration of time spent avoiding is detected with the threshold method. (See SI Table 2 for statistical details). The top two plots represent duration of time approaching (cyan) and the bottom two represent the duration of time avoiding (purple). The plots on the left are the durations measured by a human-rater and the plots on the right are from the threshold method. Each is compared between 0 and 7 days of food-deprivation.

### Supplemental Table 2

|  |  | n | statistic | df | adjusted p-value | significance |
| --- | --- | --- | --- | --- | --- | --- |
| Approach | BORIS | 54 | -2.33 | 53 | 0.0952 | ns |
| Approach | Threshold | 54 | -0.99 | 53 | 1 | ns |
| Avoid | BORIS | 54 | -0.14 | 53 | 1 | ns |
| Avoid | Threshold | 54 | 3.48 | 53 | 0.00408 | ** |

SI Table 2: Only avoidance showed a statistical difference for the threshold based quantification method that was not present in the human-rated (BORIS) data. Table shows the results of paired t-tests to compare the duration of time approaching or avoiding between 0 and 7-days of food-deprivation as measured by the two quantification methods.

### Supplemental Figure 3

SI Figure 3: Repeated measures had no consistent effect on the statistical comparisons of 0- and 7-day food-deprived *Berghia*. A) Duration of Time feeding, from top to bottom: *i.*duration of time spent feeding for animals tested 0-days food-deprived 7-days apart, *ii.* Duration of time feeding for animals tested 7-days foo-deprived, 7-days apart, and *iii.* Comparison of sated and foo-deprived measures when averaged across the two measurements for each animal. B-C) The same plots as in A) but for B) Average distance from prey and C) average heading angle with respect to prey.

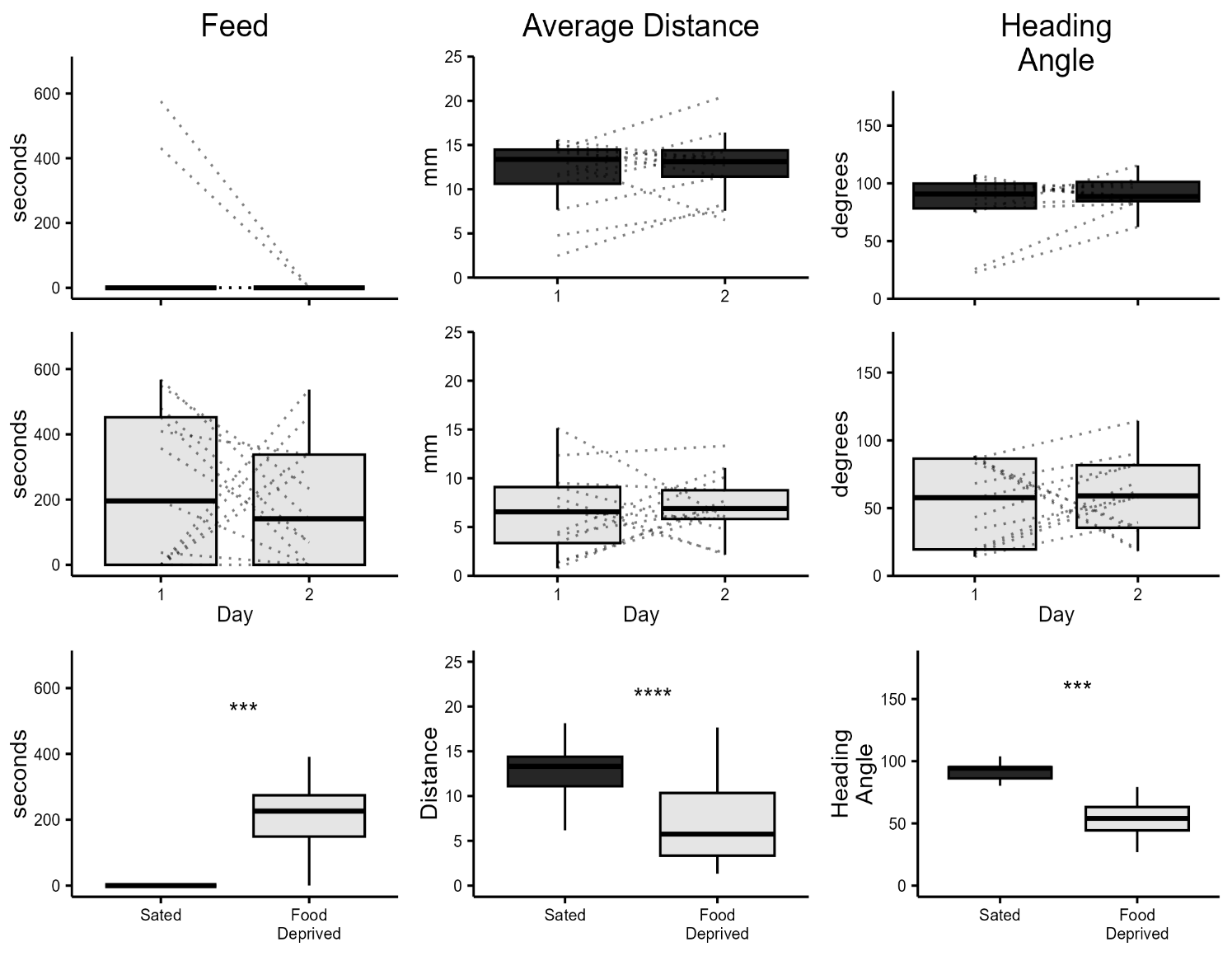

A i

B i

C i

ii

ii

ii

iii

iii

iii

### Supplemental Figure 4

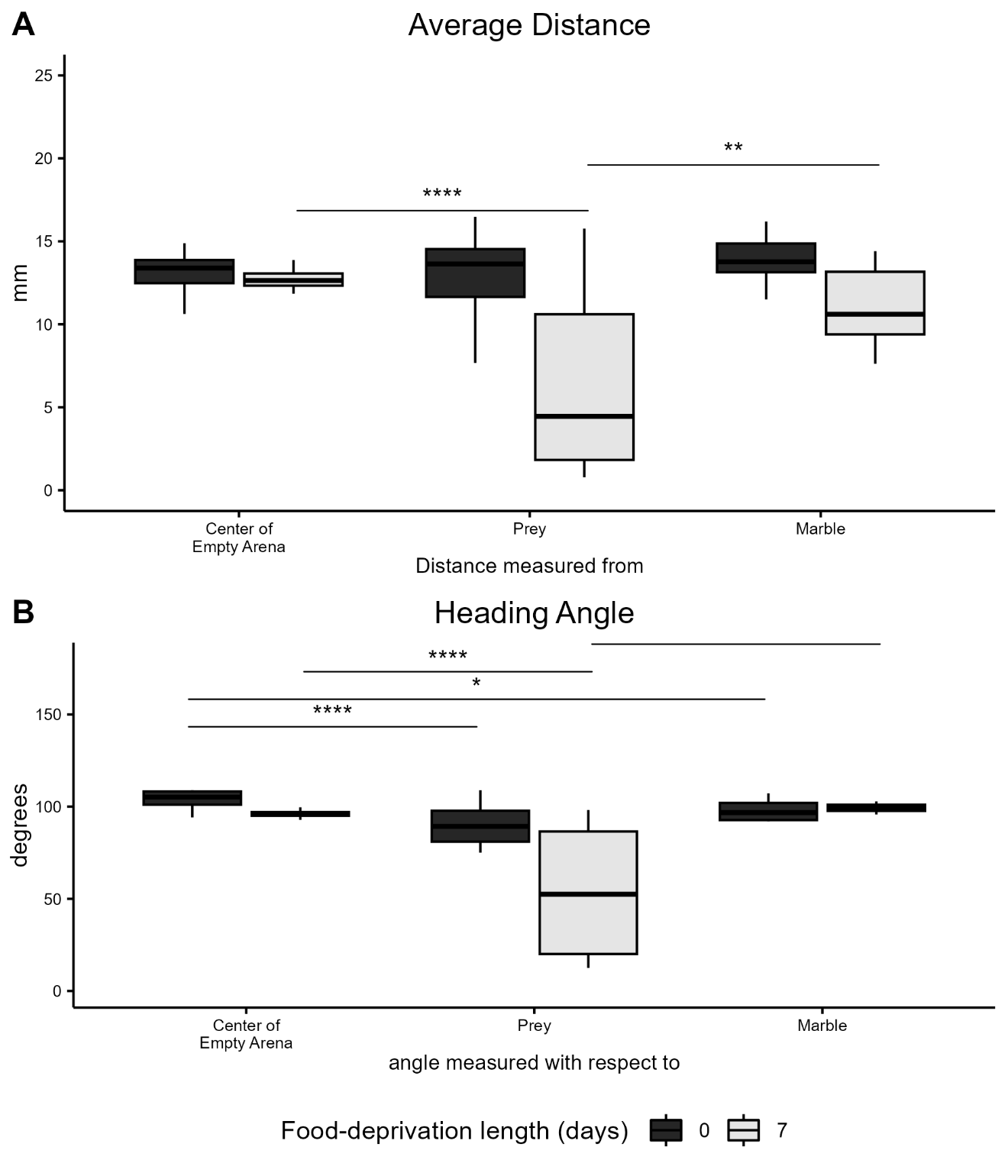

SI Fig 4: The effect of food-deprivation on distance and heading angle from prey is not explained by attraction to the center of the arena or to a neutral object (opaque white marble). A: The top plot displays the average distance from the center of an empty arena (left), from prey (middle) and from a marble paced in the center of the arena (right). There is a significant effect of the arena type on the distance measured. B: The bottom plot displays the heading angle measured with respect to the center of an empty arena (left), the prey (middle) and the marble (right). The heading angle is smaller, meaning the slug is spending more time oriented towards the prey for both sated and food-deprived animals compared to animals in an empty arena or with a marble.

### Supplemental Table 3

| Trajectory feature | Effect | DFn | DFd | F | Adjusted p-value | significance |
| --- | --- | --- | --- | --- | --- | --- |
| Emax | FDL | 1 | 185 | 1.47 | 1 | ns |
| Emax | Exp Type | 2 | 185 | 6.33 | 0.042 | * |
| Emax | FDL: Exp Type | 2 | 185 | 2.11 | 1 | ns |
| SD_speed | FDL | 1 | 185 | 0.28 | 1 | ns |
| SD_speed | Exp Type | 2 | 185 | 0.07 | 1 | ns |
| SD_speed | FDL: Exp Type | 2 | 185 | 1.24 | 1 | ns |
| distance | FDL | 1 | 185 | 2.40 | 1 | ns |
| distance | Exp Type | 2 | 185 | 9.02 | 0.003843 | ** |
| distance | FDL: Exp Type | 2 | 185 | 3.76 | 0.525 | ns |
| mean_dirchange | FDL | 1 | 185 | 0.4 | 1 | ns |
| mean_dirchange | Exp Type | 2 | 185 | 7.84 | 0.01134 | * |
| mean_dirchange | FDL: Exp Type | 2 | 185 | 2.11 | 1 | ns |
| mean_speed | FDL | 1 | 185 | 0.77 | 1 | ns |
| mean_speed | Exp Type | 2 | 185 | 4.33 | 0.294 | ns |
| mean_speed | FDL: Exp Type | 2 | 185 | 2.91 | 1 | ns |
| sd_dirchange | FDL | 1 | 185 | 0.95 | 1 | ns |
| sd_dirchange | Exp Type | 2 | 185 | 1.05 | 1 | ns |
| sd_dirchange | FDL: Exp Type | 2 | 185 | 2.84 | 1 | ns |
| sinuosity | FDL | 1 | 185 | 4.05 | 0.966 | ns |
| sinuosity | Exp Type | 2 | 185 | 4.08 | 0.399 | ns |
| sinuosity | FDL: Exp Type | 2 | 185 | 1.22 | 1 | ns |

SI Table 3: Trajectory Features of empty arena, arena with prey and arena with a marble. We analyzed features of their trajectories within the arenas including straightness (Emax), distance traveled (distance), mean and standard deviation of directional change (mean_direchange and sd_dirchange, respectively), sinuosity, mean speed and standard deviation of speed change. The trajectory features were compared to test for an effect of food-deprivation length (FDL), the experimental arena (Exp Type: empty arena, arena with prey or arena with marble), or an interaction between the two (FDL : Exp Type).

### Supplemental Figure 5

SI Fig 4: Experiment type impacted the straightness, distance travelled and directional change of trajectories. For all plots the dark grey represents sated animals and the light grey is 7-day food-deprived animals. A: Plot showing the effect of experiment type on the straightness of trajectories, where animals in the marble arena had straighter paths than animals in the other two arenas. B: Plot showing the distance travelled in the arena where animals with their prey traveled less distance than animals in an empty arena or with a marble. C: Similar pattern to B, except showing that animals with their prey showed more directional change in their trajectories.

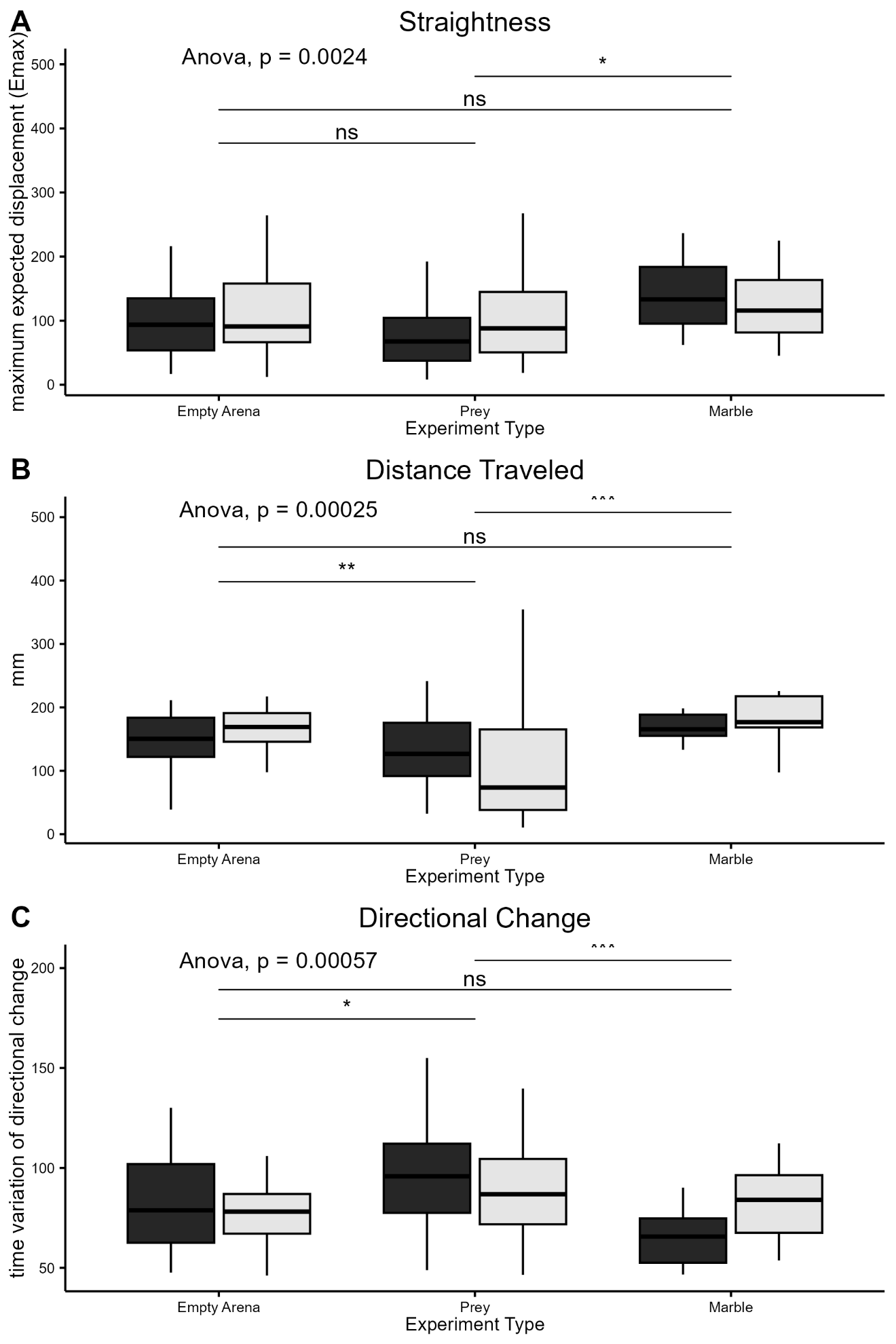

### Supplemental Figure 6

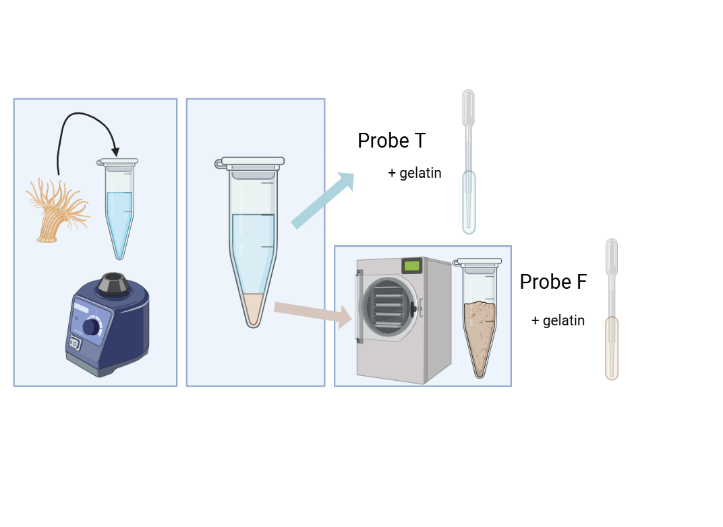

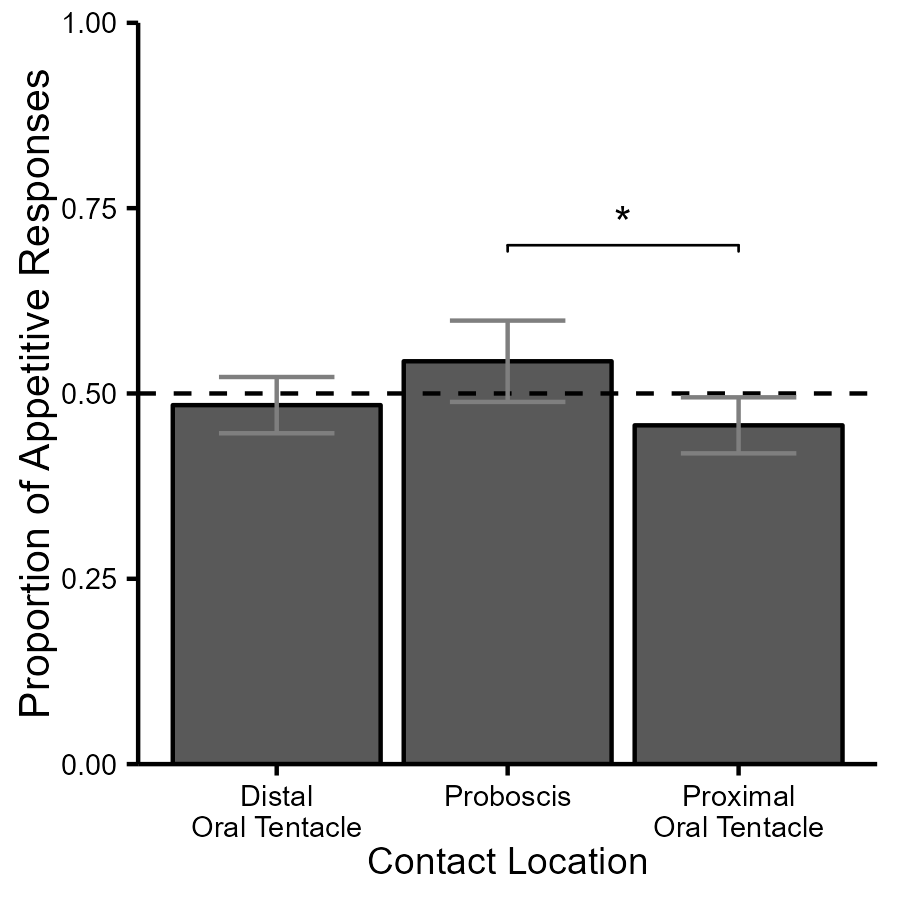

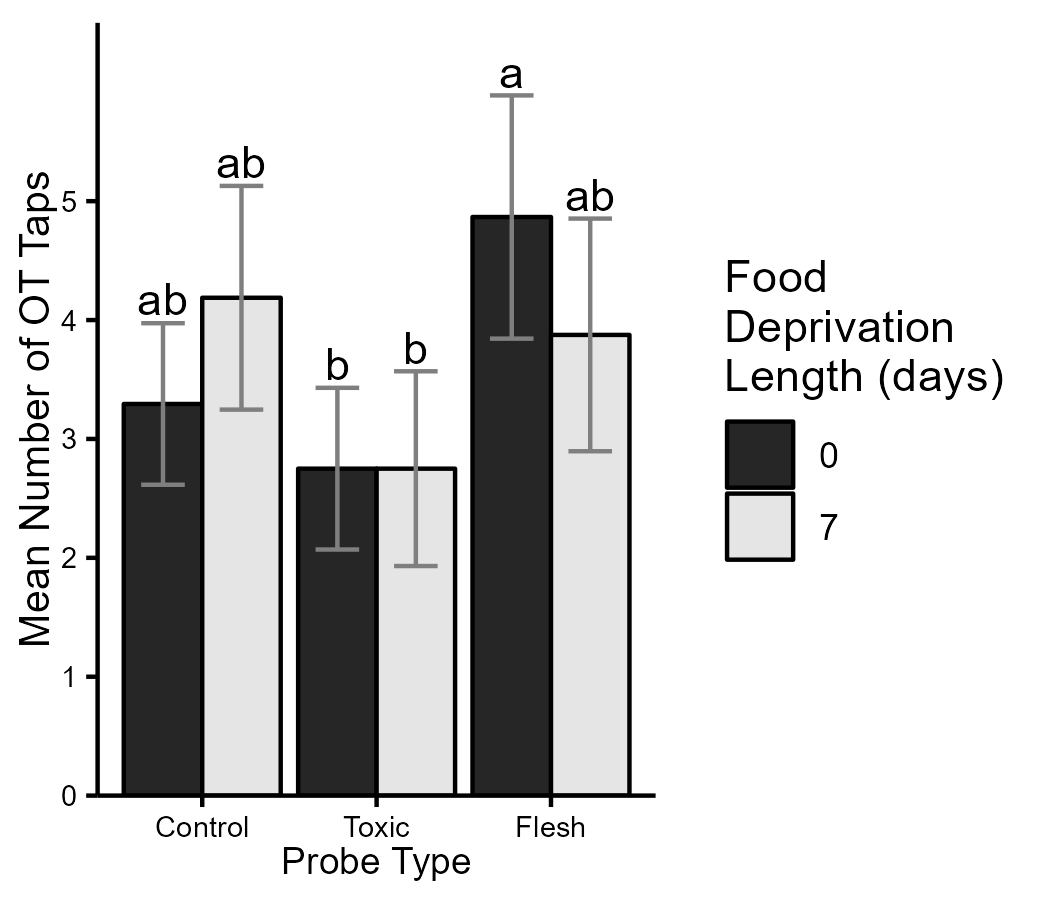

A

B

C

D

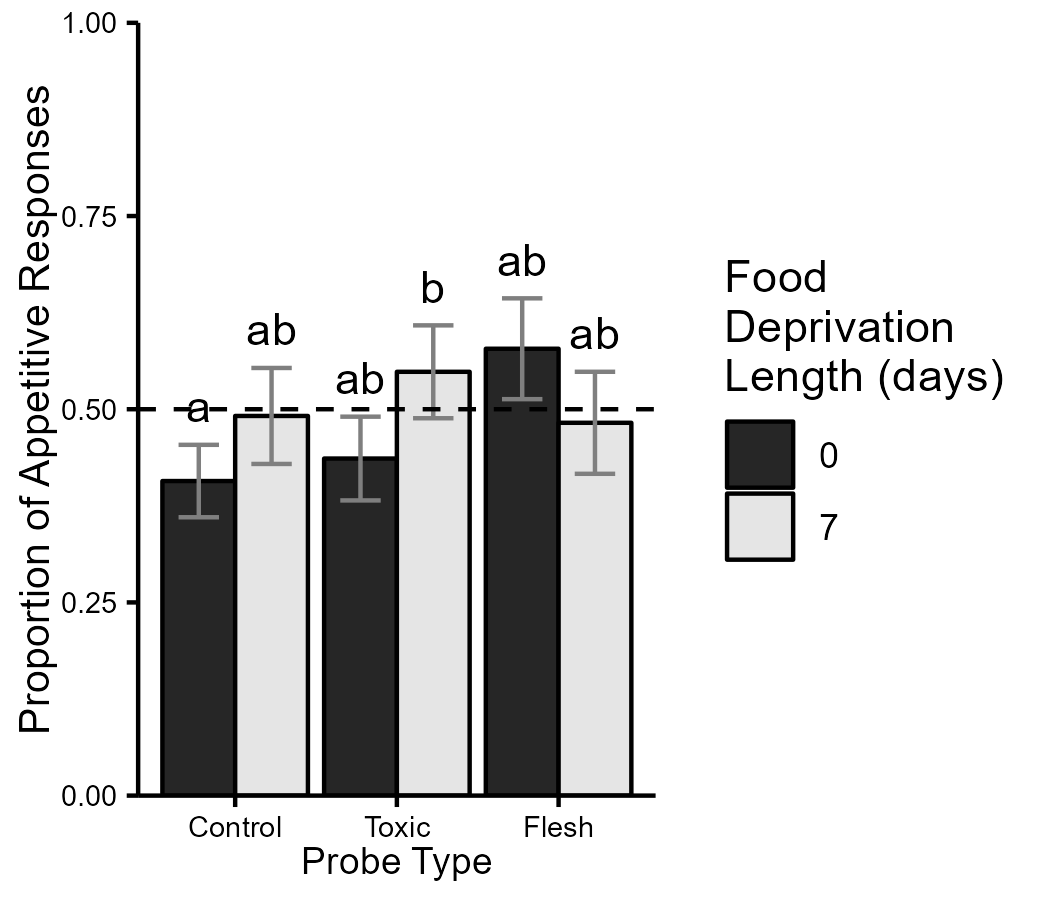

SI Figure 6: Components of prey stimuli. A) Schematic showing the process of creating probes for Experiment 4. From left to right: anemones are placed in a microcentrifuge tube with asw and vortexed, the liquid is removed and used to make the gelatin mixture for the toxin probes, the anemone is freeze dried to make a powder that is mixed with gelatin for the flesh probes. B) The proportion of responses that were appetitive in response to contact with each probe type. Dark bars represent sated animals and light bars represent food-deprived animals. A dashed line shows 0.5. Letters above the standard error error bars represent the compact letter display for the GLMM. C) Plot showing the mean number of OT taps to each type of probe during the trial. Colors, error bars and cld match B). D) Plot showing the proportion of appetitive responses in response to contact with each body part aggregated across probe type. Error bars are the standard error of the mean and dashed line is 0.5.

### Supplemental Figure 7

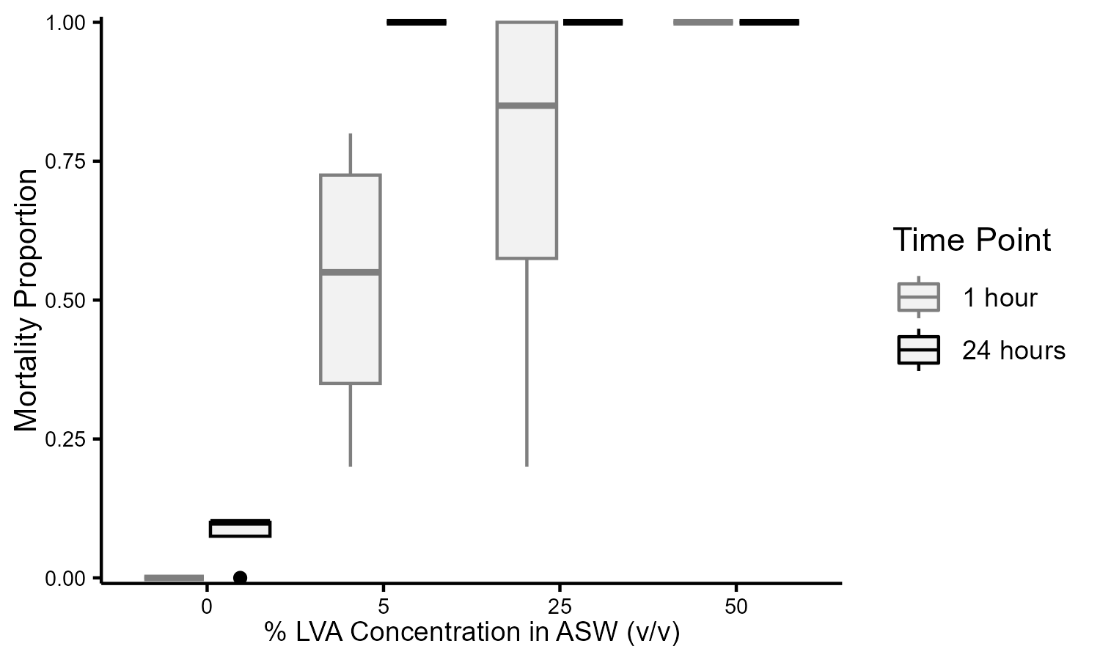

SI Figure 7: Plot showing the proportion of *Artemia* that died for each concentration of anemone toxin from a vortexed anemone (%LVA) in asw for two time points (1 hour (light) and 24 hours (dark).

### Supplemental Table 4

SI Table 4: Experiment 4 model output.

| Generalized Linear Model (GLM) | | | | |
| --- | --- | --- | --- | --- |
|  | Estimate | Standard Error | Z value | Pr(>\|z\|) |
| (Intercept) | 0.3924 | 0.114 | 3.436 | 0.000591 |
| Toxic Probe | -0.1229 | 0.1502 | -0.818 | 0.413325 |
| Flesh Probe | 0.9465 | 0.1552 | 6.098 | 1.08e-09 |
| Food-deprivation Length (7 days) | 0.1047 | 0.1439 | 0.727 | 0.466948 |
| Proboscis contact | -0.2502 | 0.1375 | -1.820 | 0.068829 |
| Proximal Oral Tentacle contact | -0.4188 | 0.1034 | -4.052 | 5.08e-05 |
| Interaction:  Toxic probe: Food-deprivation Length (7 days) | 0.4465 | 0.2283 | 1.955 | 0.050540 |
| Interaction:  Flesh probe: Food-deprivation Length (7 days) | -0.9147 | 0.2278 | -4.015 | 5.94e-05 |

### Supplemental Table 5

BORIS ethogram used for Experiments 1, 2 and 3.

| Behavior code | Behavior type | Description | Key | Excluded behaviors | Modifiers |
| --- | --- | --- | --- | --- | --- |
| Explore | State event | Movement that is not oriented towards or directly away from the prey item heading angle | e | Approach,Avoid,Roll-Over | |
| Approach | State event | Movement that is oriented towards the prey | a | Explore,Avoid,Feed,Roll-Over | |
| Protrude | State event | Berghia everts its buccal mass aka proboscis an m shaped structure appears in front of their snut | p | Feed,Roll-Over | |
| Avoid | State event | Movement that is oriented away from the prey | v | Explore,Approach,Feed,Roll-Over | |
| Feed | State event | Berghia is in direct contact with the prey and is swallowing the prey | n | Approach,Protrude,Avoid,Roll-Over | |
| Tail Withdrawal | Point event | Berghia retracts its tail such that the tail disappears from view or retracts its posterior body such that its length shortens with movement only occurring in the posterior body | t |  |  |
| Oral Tentacle Withdrawal | Point event | Berghia retracts one of its oral tentacles flexing dorsally at the proximal end oral tentacle appears to shorten in length | q |  |  |
| Head Withdrawal | Point event | Berghia retracts its head towards the middle of its body cerata perform a rhinal bristle eyes and head spot disappear from view tail and posterior body remain in place | h |  |  |
| Oral Tentacle Tap | Point event | Distal end of the oral tentacle makes contact with a stimulus and briefly flexes away from the stimulus often dorsally and then immediately returns to a resting position or makes contact again | d |  |  |
| J-Turn | Point event | | j |  |  |
| F-Turn | Point event | | f |  |  |
| Scrunch | Point event | | s |  |  |
| Bite | Point event | Berghia attempts to bite a stimulus buccal mass can be seen moving outward and then inward | b |  |  |
| Head Wave | Point event | With posterior body stationary Berghia bends at the mid body and moves its head laterally and back to midline in both left and right directions | z |  |  |
| 360-Turn | Point event | Berghia turns 360 degrees or more in one direction entire body should complete a circle | 3 |  |  |
| Pause | Point event | | u |  |  |
| Rhinal Bristle | Point event | Berghias first cerata bunch become erect and move medially and towards the head and rhinophores often occluding the head and eyes | k |  |  |
| Lateral Bristle | Point event | | l |  |  |
| Medial Bristle | Point event | | m |  |  |
| 180-Turn | Point event | Berghia lifts its head from the substrate and rotates its head around the middle of its body until it is pointed away from the anemone returns head to the substrate and move in the new direction with the posterior body following | 8 |  |  |
| Curl | Point event | Berghia laterally flexes its body at the level of the middle of its body until its head is parallel to its posterior body and pauses in this C shape tail and posterior body remain in place | c |  |  |
| Rear | Point event | Berghia lifts its head dorsally off the substrate for more than 0.5 seconds | x |  |  |
| Orient | Point event | From a neutral heading to their prey or from a directionally opposite heading to their prey Berghia turns its head towards their prey and proceeds in that direction with their body following | o |  |  |
| Oral Tentacle W Pose | Point event | Both of Berghias oral tentacles flex dorsolaterally at the proximal end such that the tips end up level with the head and for a W shape this pose is held for at least 0.5 seconds | w |  |  |
| Cross Oral Tentacles | Point event | Both of Berghias oral tentacles flex dorsomedially at the proximal end such that the tips end up level with the head and in line with the body often they will form an X shape and the tips will cross midline | r |  |  |
| AnemoneStuck | State event | | 7 |  |  |
| Tentacle Withdrawal | Point event | Following a stimulus the anemone rapidly withdraws its tentacle towards its mouth-anus will often seem to deflate and shrink | 6 |  |  |
| Body Column Scrunch | Point event | Body column of the anemone rapidly shortens in length | 9 |  |  |
| Acontia Ejection | Point event | Anemone ejects its acontia white thin strands can be seen coiling and uncoiling near the base of the anemone | 0 |  |  |
| Roll-Over | State event | | 4 | Explore,Approach,Protrude,Avoid,Feed | |
| Mouth Popping | Point event | | 1 |  |  |
| Tentacle_snut | Point event | | g |  |  |
| Tentacle_head | Point event | | i |  |  |
| Tentacle_distalOT | Point event | | y |  |  |
| Tentacle_proximalOT | Point event | | 2 |  |  |
| Tentacle_body | Point event | | 5 |  |  |
| Tentacle_proboscis | Point event | | , |  |  |

### Supplemental Table 6

BORIS ethogram used for experiment 4:

| Behavior code | Behavior type | Description | Key | Modifiers |
| --- | --- | --- | --- | --- |
| probe_contact | Point event | | c | location:proboscis (1),distal_oral_tentacle (2),proximal_oral_tentacle (3),middle_oral_tentacle (4),lips (5),head (6) |
| probe_location | State event | | l | location:left_lateral (1),right_lateral (2),medial (3) |
| protrude | State event | | p |  |
| ignore | State event | | i |  |
| rear | Point event | | r |  |
| appetitive | Point event | | a | appetitive behaviors:bite (1),OT tap (2),cross OT (3),turn towards (4) |
| aversive | Point event | | v | aversive behaviors:bristle (1),OT_withdrawal (2),scrunch (3),head withdrawal (4),turn away (5) |
| OT_W_shape | Point event | | w |  |
